# Stochastic Yield Catastrophes and Robustness in Self-Assembly

**DOI:** 10.1101/660340

**Authors:** Florian M. Gartner, Isabella R. Graf, Patrick Wilke, Philipp M. Geiger, Erwin Frey

## Abstract

A guiding principle in self-assembly is that, for high production yield, nucleation of structures must be significantly slower than their growth. However, details of the mechanism that impedes nucleation are broadly considered irrelevant. Here, we analyze self-assembly into finite-sized target structures employing mathematical modeling. We investigate two key scenarios to delay nucleation: (i) by introducing a slow activation step for the assembling constituents and, (ii) by decreasing the dimerization rate. These scenarios have widely different characteristics. While the dimerization scenario exhibits robust behavior, the activation scenario is highly sensitive to demographic fluctuations. These demographic fluctuations ultimately disfavor growth compared to nucleation and can suppress yield completely. The occurrence of this stochastic yield catastrophe does not depend on model details but is generic as soon as number fluctuations between constituents are taken into account. On a broader perspective, our results reveal that stochasticity is an important limiting factor for self-assembly and that the specific implementation of the nucleation process plays a significant role in determining the yield.

## 1 Introduction

Efficient and accurate assembly of macromolecular structures is vital for living organisms. Not only must resource use be carefully controlled, but malfunctioning aggregates can also pose a substantial threat to the organism itself ^1, 2^. Furthermore, artificial self-assembly processes have important applications in a variety of research areas like nanotechnology, biology, and medicine ^3–5^. In these areas, we find a broad range of assembly schemes. While viral capsids are generally assembled from identical substructures ^6–8^, DNA-brick-based assembly is used to artificially construct almost any desired configuration starting from large numbers of different constituents ^9–12^. Notwithstanding these differences, a generic self-assembly process always includes three key steps: First, subunits must be made available, e.g. by gene expression, or rendered competent for binding, e.g. by nucleotide exchange ^13, 14^ (‘activation’). Second, the formation of a structure must be initiated by a nucleation event (‘nucleation’). Due to cooperative or allosteric effects in binding, there might be a significant nucleation barrier ^15–19^. Third, following nucleation, structures grow via aggregation of substructures (‘growth’). To avoid kinetic traps that may occur due to irreversibility or very slow disassembly of substructures ^20, 21^, structure nucleation must be significantly slower than growth ^6, 9, 10, 22–24^. In this manuscript we investigate, for a given target structure, whether the nature of the specific mechanism employed in order to slow down nucleation influences the yield of assembled product. To address this question, we examine a generic model that incorporates the key elements of self-assembly outlined above.

## 2 Model definition

We consider a set of *S* different species of constituents denoted by 1, …, *S* which assemble into rings of size *L*. The cases *S* = 1 and 1 < *S* ≤ *L* (*S* = *L*) are denoted as homogeneous and partially (fully) heterogeneous, respectively. The homogeneous model builds on previous work on virus capsid and linear protein filament assembly ^15, 20, 25–27^. The heterogeneous model in turn links to previous model systems used to study, for example, DNA-brick-based assembly of heterogeneous structures ^28–30^. We emphasize that, even though strikingly similar experimental realizations of our model exist ^11, 12, 31^, it is not intended to describe any particular system. The ring structure represents a general linear assembly process involving building blocks with equivalent binding properties and resulting in a target of finite size. In order to test the robustness of our findings, we furthermore perform stochastic simulations of systems with non-linear assembly paths and no translational symmetry (see Figs. 5 and 6).

The assembly process starts with *N* inactive monomers of each species. We use *C* = *N/V* to denote the initial concentration of each monomer species, where *V* is the reaction volume. Monomers are activated independently at the same per capita rate *α*, and, once active, are available for binding. Binding takes place between constituents of species with periodically consecutive indices, for example 1 and 2 or *S* and 1 (Fig. 1). To avoid ambiguity, we restrict ring sizes to integer multiples of the number of species *S*. Polymers, i.e., incomplete ring structures, grow via consecutive attachment of monomers. For simplicity, polymer-polymer binding is disregarded at first, as it is typically assumed to be of minor importance ^6, 15, 28, 32^. To probe the robustness of the model, later we consider an extended model including polymer-polymer binding for which the results are qualitatively the same (see Fig. 5 and the discussion). Furthermore, it has been observed that nucleation phenomena play a critical role for self-assembly processes ^9, 10, 15, 22^. So it is in general necessary to take into account a critical nucleation size, which marks the transition between slow particle nucleation and the faster subsequent structure growth ^18, 25, 28, 33^. We denote this critical nucleation size by *L*_nuc_, which in terms of classical nucleation theory corresponds to the structure size at which the free energy barrier has its maximum. For *l* < *L*_nuc_ attachment of monomers to existing structures and decay of structures into monomers take place at size-dependent reaction rates *µ*_*l*_ and *δ*_*l*_, respectively (Fig. 1). Here, we focus on identical rates *µ*_*l*_ = *µ* and *δ*_*l*_ = *δ*. A discussion of the general case is given in the Supplemental Material ^34^. Above the nucleation size, polymers grow by attachment of monomers with reaction rate *ν* ≥ *µ* per binding site. As we consider successfully nucleated structures to be stable on the observational time scales, monomer detachment from structures above the critical nucelation size is neglected ^15, 28^. Complete rings neither grow nor decay (absorbing state).

**Figure 1:**
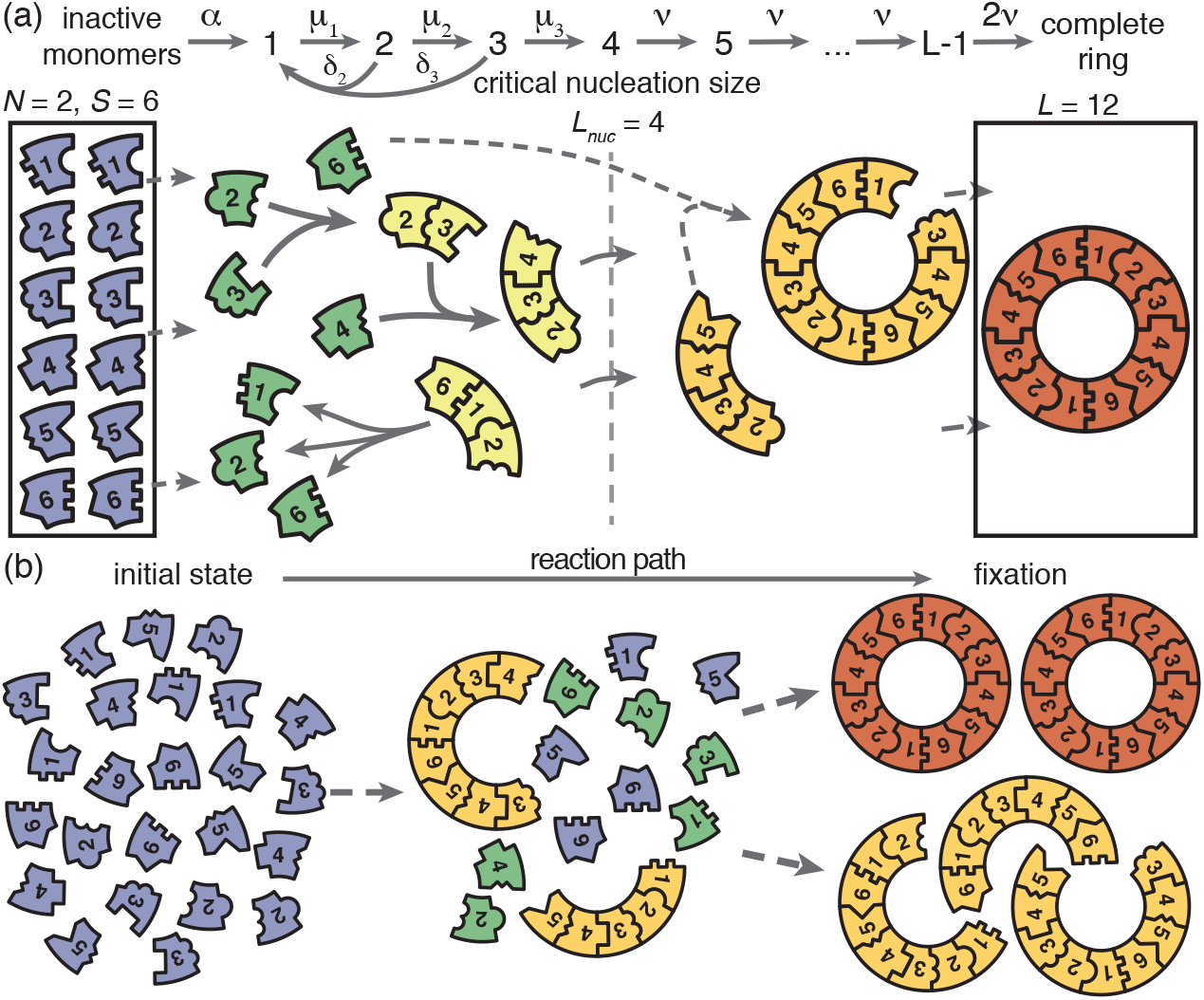
Schematic description of the model. (a) Rings of size *L* are assembled from *S* different particle species. *N* monomers of each species are initially in an inactive state and are activated at the same per-capita rate *α*. Once active, species with periodically consecutive index can bind to each other. Structures grow by attachment of single monomers. Below a critical nucleation size (*L*_nuc_), structures of size *l* grow and decay into monomers at size-dependent rates *µ_l_* and *δ*_*l*_, respectively. Above the critical size, polymers are stable and grow at size-independent rate *ν* until the ring is complete (the absorbing state). (b) Illustration of depletion traps. If nucleation is slow compared to growth, initiated structures are likely to be completed. Otherwise, many stable nuclei will form that cannot be completed before resources run out.

We investigate two scenarios for the control of nucleation speed, first separately and then in combination. For the ‘activation scenario’ we set *µ* = *ν* (all binding rates are equal) and control the assembly process by varying the activation rate *α*. For the ‘dimerization scenario’ all particles are inherently active (*α* → *∞*) and we control the assembly process by varying the dimerization rate *µ* (we focus on *L*_nuc_ = 2). We quantify the quality of the assembly process in terms of the assembly yield, defined as the number of successfully assembled ring structures relative to the maximal possible number *NS/L*. Yield is measured when all resources have been used up and the system has reached its final state. We simulated our system both stochastically via Gillespie’s algorithm ^35^ and deterministically as a set of ordinary differential equations corresponding to chemical rate equations (see Supplemental Material ^34^).

## 3 Results

### Deterministic behavior in the macroscopic limit

First, we consider the macroscopic limit, *N* ≫ 1, and investigate how assembly yield depends on the activation rate *α* (activation scenario) and the dimerization rate *µ* (dimerization scenario) for *L*_nuc_ = 2. Here, the deterministic description coincides with the stochastic simulations (Fig. 2(a) and (b)). For both high activation and high dimerization rates, yield is very poor. Upon decreasing either the activation rate (Fig. 2(a)) or the dimerization rate (Fig. 2(b)), however, we find a threshold value, *α*_th_ or *µ*_th_, below which a rapid transition to the perfect yield of 1 is observed both in the deterministic and stochastic simulation. By exploiting the symmetries of the system with respect to relabeling of species, one can show that, in the deterministic limit, the behavior is independent of the number of species *S* (for fixed *L* and *N*, see Supplemental Material ^34^). Consequently, all systems behave equivalently to the homogeneous system and yield becomes independent of *S* in this limit. Note, however, that equivalent systems with differing *S* have different total numbers of particles *SN* and hence assemble different total numbers of rings.

**Figure 2:**
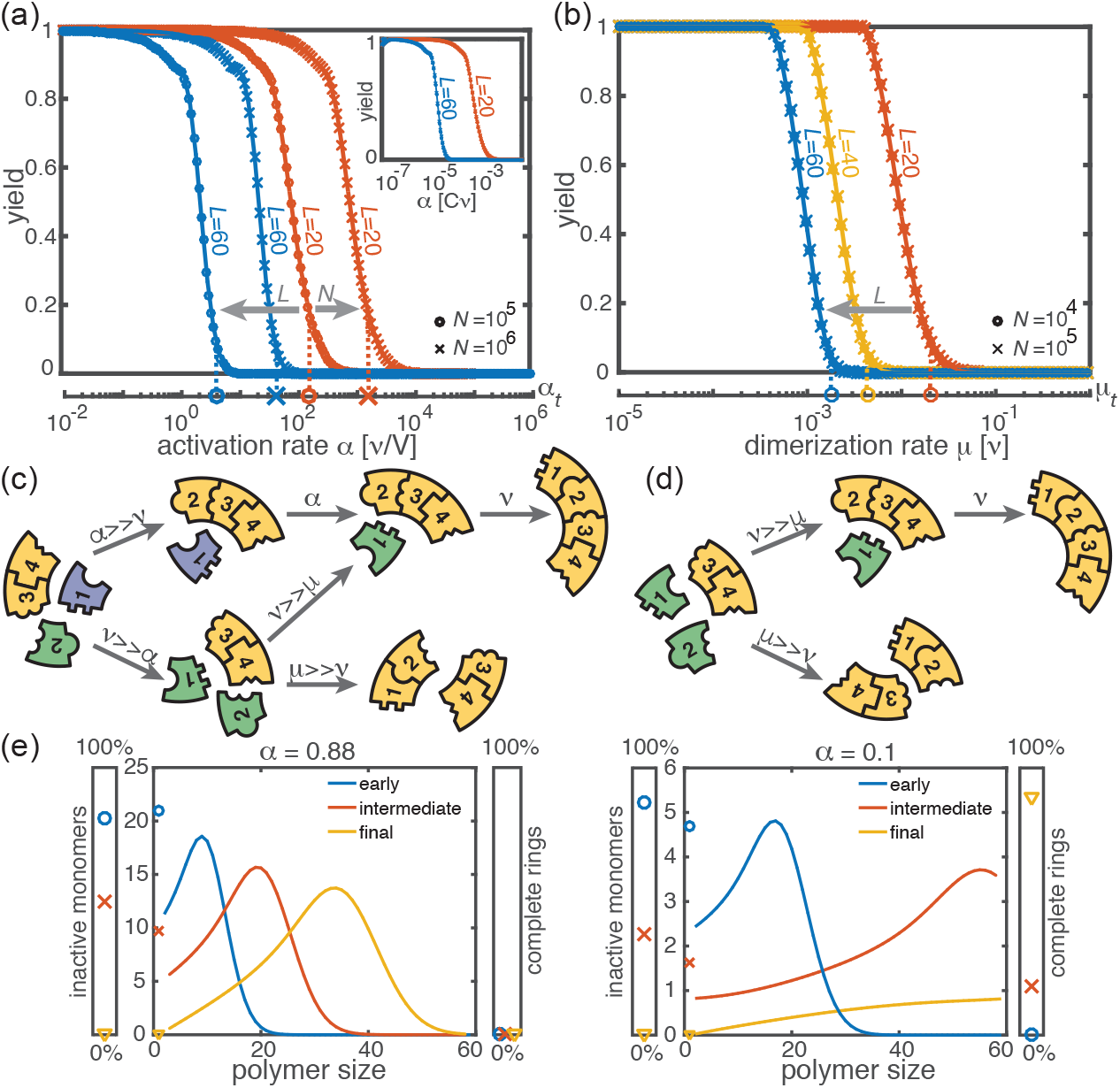
Deterministic behavior in the macroscopic limit *N* ≫ 1. (a, b) Yield for different particle numbers *N* (symbols) and ring sizes *L* (colors) for *L*_nuc_ = 2. Decreasing either (a) the activation rate (‘activation scenario’: *µ* = *ν*) or (b) the dimerization rate (‘dimerization scenario’: *α* → *∞*) achieves perfect yield. The stochastic simulation results (symbols) agree exactly with the integration of the chemical rate equations (lines). The threshold values (Eq. 1) are indicated by the vertical dashed lines. Plotting yield against the dimensionless quantity *α/*(*νC*) causes the curves for different *C* to collapse into a single master curve (inset in a). For both scenarios there is no dependency on the number of species *S* in the deterministic limit. (c, d) Illustration showing how depletion traps are avoided by either slow activation (c) or slow dimerization (d). If the activation or the dimerization rate is small (large) compared to the growth rate, assembly paths leading to complete rings are favored (disfavored). (e) Deterministically, the size distribution of polymers behaves like a wave, and is shown for large and small activation rate for *L* = 60, *L*_nuc_ = 2, *N* = 10000 and *µ* = *ν* = 1. The distributions are plotted for early, intermediate and final simulation times. The respective percentage of inactive monomers and complete rings is indicated by the symbols in the scale bar on the left or right.

Decreasing the activation rate reduces the concentration of active monomers in the system. Hence growth of the polymers is favored over nucleation, because growth depends linearly on the concentration of active monomers while nucleation shows a quadratic dependence. Likewise, lower dimerization rates slow down nucleation relative to growth. Both mechanisms therefore restrict the number of nucleation events, and ensure that initiated structures can be completed before resources become depleted (see Fig. 2(c) and (d)).

Mathematically, the deterministic time evolution of the polymer size distribution *c*(*l, t*) is described by an advection-diffusion equation ^36, 37^ with advection and diffusion coefficients depending on the instantaneous concentration of active monomers (see Supplemental Material ^34^). Solving this equation results in the wavefront of the size distribution advancing from small to large polymer sizes (Fig. 2(e)). Yield production sets in as soon as the distance travelled by this wavefront reaches the maximal ring size *L*. Exploiting this condition, we find that in the deterministic system for *L*_nuc_ = 2, a non-zero yield is obtained if either the activation rate or the dimerization rate remains below a corresponding threshold value, i.e. if *α* < *α*_th_ or *µ* < *µ*_th_, where

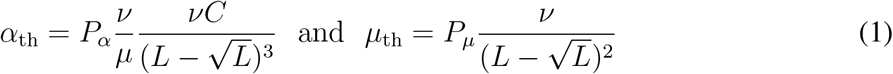

(see Supplemental Material ^34^) with proportionality constants 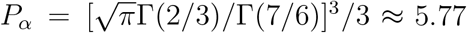 and *P*_*µ*_ = *π*^2^/2 ≈ 4.93. These relations generalize previous results ^33^ to finite activation rates, and highlight important differences between the two scenarios (where *α* → *∞* and *µ* = *ν*, respectively): While *α*_th_ decreases cubically with the ring size *L*, *µ*_th_ does so only quadratically. Furthermore, the threshold activation rate *α*_th_ increases with the initial monomer concentration *C* (Fig. 2(a)). Consequently, for fixed activation rate, the yield can be optimized by increasing *C*. In contrast, the threshold dimerization rate is independent of *C* and the yield curves coincide for *N* ≫ 1. Finally, if *α* is finite and *µ < ν*, the interplay between the two slow-nucleation scenarios may lead to enhanced yield. This is reflected by the factor *ν/µ* in *α*_th_, and we will come back to this point later when we discuss the stochastic effects.

In summary, for large particle numbers (*N ≫* 1), perfect yield can be achieved in two different ways, independently of the heterogeneity of the system -by decreasing either the activation rate (activation scenario) or the dimerization rate (dimerization scenario) below its respective threshold value.

### Stochastic effects in the case of reduced resources

Next, we consider the limit where the particle number becomes relevant for the physics of the system. In the activation scenario, we find a markedly different phenomenology if resources are sparse. Figure 3(a) shows the dependence of the yield on the activation rate for different, low particle numbers in the completely heterogeneous case (*S* = *L*). Whereas the deterministic theory predicts perfect yield for small activation rates, in the stochastic simulation yield saturates at an imperfect value *y*_max_ < 1. Reducing the particle number *N* decreases this saturation value *y*_max_ until no finished structures can be produced, an effect we term ‘stochastic yield catastrophe’. Moreover, *y*_max_ increases with decreasing number of species *S* (Fig. 3(c)). For the fully homogeneous case, *S* = 1, a perfect yield of 1 is always achieved for *α →* 0. Finally, increasing the nucleation size *L*_nuc_, and with it the reversibility of binding, also increases *y*_max_. This indicates that both heterogeneity of the target structure and irreversibility of binding make the system more susceptible to stochastic effects. The stochastic yield catastrophe is mainly attributable to fluctuations in the number of active monomers. In the deterministic (mean-field) equation the different particle species evolve in balanced stoichiometric concentrations. However, if activation is much slower than binding, the number of active monomers present at any given time is small, and the mean-field assumption of equal concentrations is violated due to fluctuations (*S >* 1). Activated monomers then might not fit any of the existing larger structures and would instead initiate new structures. Due to the effective enhancement of nucleation rate the resulting polymer size distribution has a higher amplitude than that predicted deterministically (Fig. 3(d)) and the system is prone to depletion traps. A similar broadening of the size distribution has been reported in the context of stochastic coagulation-fragmentation of identical particles ^38^.

**Figure 3:**
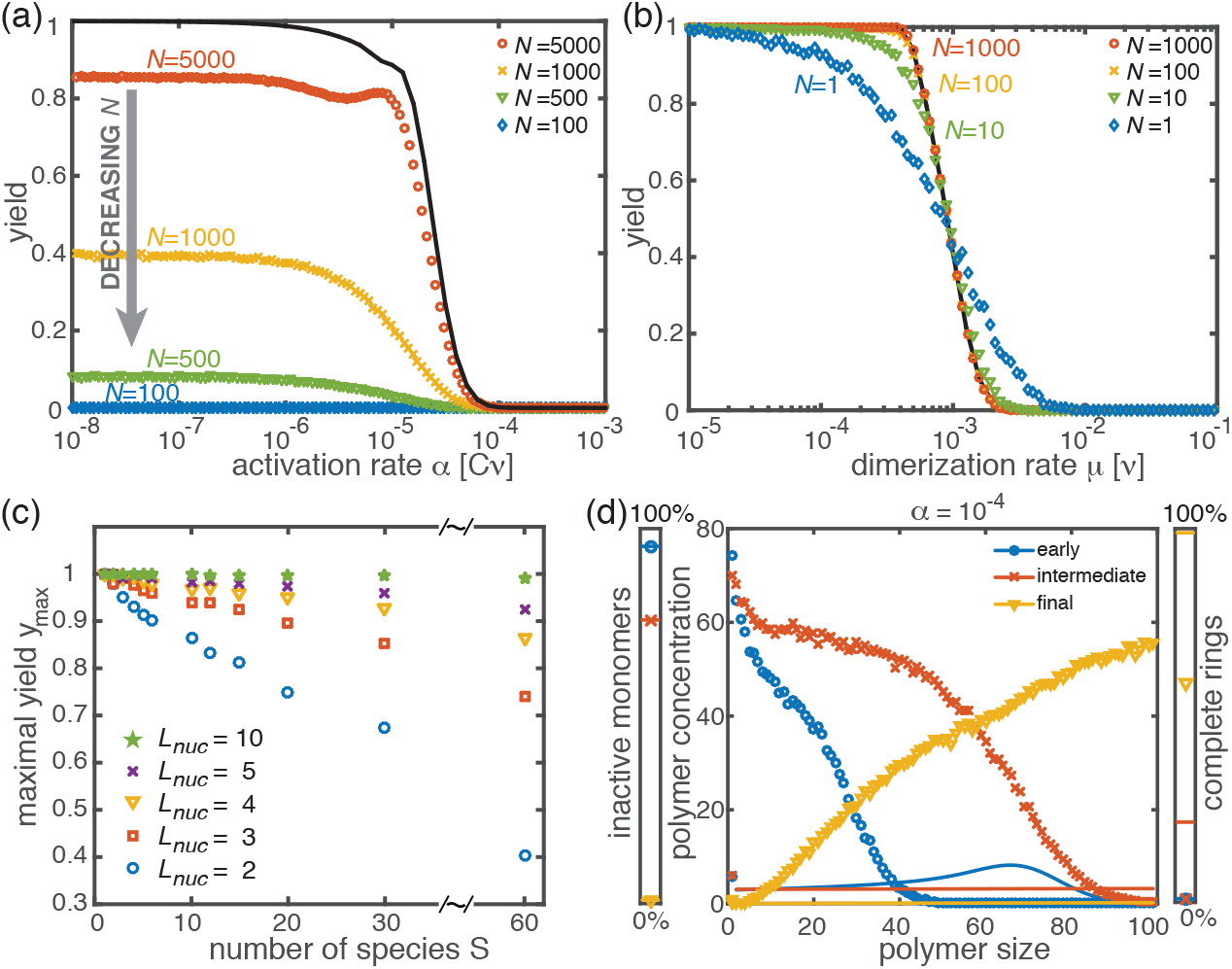
Stochastic effects in the case of reduced resources. (a, b) Yield of the fully heterogeneous system (*S* = *L*) for reduced number of particles (symbols) for *L* = 60 and *L*_nuc_ = 2. In the activation scenario, at low activation rates the yield saturates at an imperfect value *y*_max_, which decreases with the number of particles (a). This finding disagrees with the deterministic prediction of perfect yield for *α* → 0. In contrast, the dimerization scenario robustly exhibits the maximal yield of 1 for small *N* (b). (c) Dependence of maximal yield in the activation scenario on the number of species *S* for *L* = 60. Symbols indicate different values of the critical nucleation size *L*_nuc_. The total number of particles is held constant at *SN* = 60000. The impact of stochastic effects strongly depends on the number of species. The homogeneous system is not subject to stochastic effects at all. Higher reversibility for larger *L*_nuc_ also mitigates stochastic effects. (d) Polymer size distribution for the activation scenario (symbols) in comparison with its deterministic prediction (lines) for *S* = *L* = 100, *N* = 1000 and *L*_nuc_ = 2. Due to the enhanced number of nucleation events, the stochastic wave encompasses far more structures and moves more slowly. As a result, it does not quite reach the absorbing boundary.

In the dimerization scenario, in contrast, there is no stochastic activation step. All particles are available for binding from the outset. Consequently, stochastic effects do not play an essential role in the dimerization scenario and perfect yield can be reached robustly for all system sizes, regardless of the number of species *S* (Fig. 3(b)).

### Non-monotonic yield curves for a combination of slow dimerization and activation

So far, the two implementations of the ‘slow nucleation principle’ have been investigated separately. Surprisingly, we observe counter-intuitive behavior in a mixed scenario in which both dimerization and activation occur slowly (i.e., *µ* < *ν*, *α* < *∞*). Figure 4 shows that, depending on the ratio *µ/ν*, the yield can become a non-monotonic function of *α*. In the regime where *α* is large, nucleation is dimerization-limited; therefore activation is irrelevant and the system behaves as in the dimerization scenario for *α* → *∞*. Upon decreasing *α* we then encounter a second regime, where activation and dimerization jointly limit nucleation. The yield increases due to synergism between slow dimerization and activation (see *µ/ν* dependence of *α*_th_, Eq. 1), whilst the average number of active monomers is still high and fluctuations are negligible. Finally, a stochastic yield catastrophe occurs if *α* is further reduced and activation becomes the limiting step. This decline is caused by an increase in nucleation events due to relative fluctuations in the availability of the different species (“fluctuations between species”). This contrasts the deterministic description where nucleation is always slower for smaller activation rate. Depending on the ratio *µ/ν*, the ring size *L* and the particle number *N*, maximal yield is obtained either in the dimerization-limited (red curves, Fig. 4), activation-limited (blue curve, Fig. 4(b)) or intermediate regime (green and orange curves).

**Figure 4:**
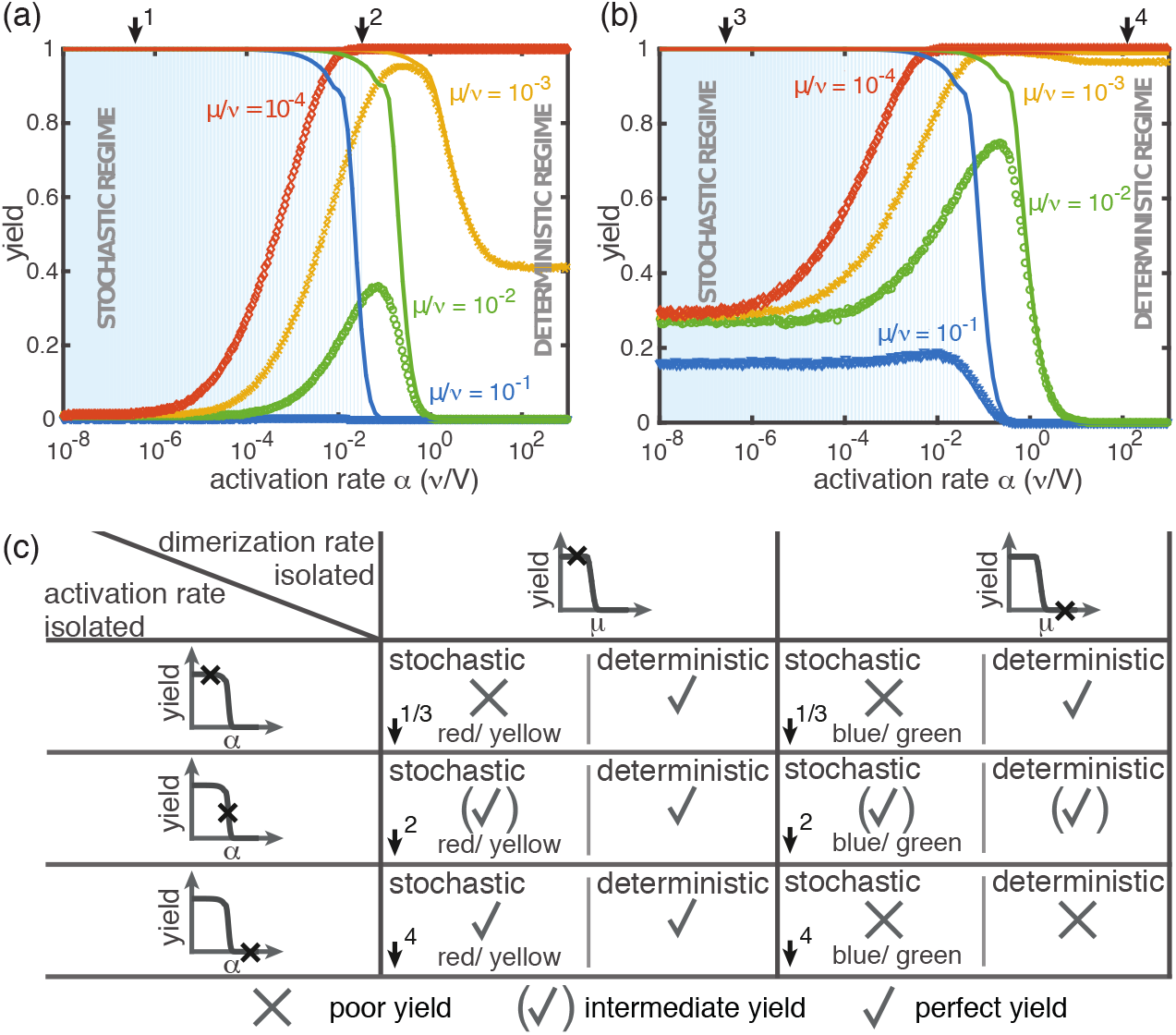
Yield for a combination of slow dimerization and activation. (a, b) Dependence of the yield of the fully heterogeneous system on the activation rate *α* for *N* = 100 and different values of the dimerization rate (colors/symbols) for *L* = 60 (a) and *L* = 40 (b). For large activation rates yield behaves deterministically (lines). In contrast, for small activation rates stochastic effects (blue shading) lead to a decrease in yield. Depending on the parameters, the yield maximum is attained in either the deterministic, stochastic or intermediate regime. (c) Table summarizing the qualitative behavior of the yield (poor/intermediate/perfect) for a combination of dimerization and activation rates for both the deterministic and the stochastic limit. The columns correspond to low and high values of the dimerization rate, as indicated by the marker in the corresponding deterministic yield curve at the top of the column. Similarly, the rows correspond to low, intermediate and high activation rates. Arrows and colors indicate where and for which curve this behavior can be observed in (a) and (b). Deviations between the deterministic and stochastic limits are most prominent for low activation rates.

### Robustness of the results to model modifications

In our model, the reason for the stochastic yield catastrophe is that - due to fluctuations between species - the effective nucleation rate is strongly enhanced. As a result, there is an excess of structures that ultimately cannot be completed. This is due to the fact that larger structures can not combine. Thus, a natural question to ask is whether relaxing the constraint that polymers cannot bind other polymers will resolve the stochastic yield catastrophe. To answer this question, we performed stochastic simulations for an extended model showing that the stochastic yield catastrophe indeed persists. In this extended model, polymer-polymer binding is taken into account in addition to growth via monomer attachment (Fig.5). In detail, we assume that two structures of arbitrary size (and with combined length ≤ *L*) bind at rate *ν* if they fit together, i.e. if the left (right) end of the first structure is periodically continued by the right (left) end of the second one. Realistically, the rate of binding between two structures is expected to decrease with the motility and thus the sizes of the structures. In order to assess the effect of polymer-polymer binding, we focus on the worst case where the rate for binding is independent of the size of both structures. If a stochastic yield catastrophe occurs for this choice of parameters, we expect it to be even more pronounced in all the “intermediate cases”. Fig. 5 shows the dependence of the yield on the activation rate in the polymer-polymer model. As before, yield increases below a critical activation rate and then saturates at an imperfect value for small activation rates. Decreasing the number of particles per species, decreases this saturation value. Compared to the original model, the stochastic yield catastrophe is mitigated but still significant: For structures of size *S* = *L* = 100, yield saturates at around 0.87 for *N* = 100 particles per species and at around 0.33 for *N* = 10 particles per species. We thus conclude that polymer-polymer binding indeed alleviates the stochastic yield catastrophe but does not resolve it. Since binding only happens between consecutive species, structures with overlapping parts intrinsically can not bind together and depletion traps occur. Taken together, also in the extended model, fluctuations in the availability of the different species lead to an excess of intermediate-sized structures that get kinetically trapped due to structural mismatches. Note that in the extreme case of *N* = 1, incomplete polymers can always combine into 1 final ring structure so that in this case yield is always 1. Analogously, for high activation rates yield is improved for *N* = 10 compared to *N* ≥ 50 (Fig. 5 b).

**Figure 5:**
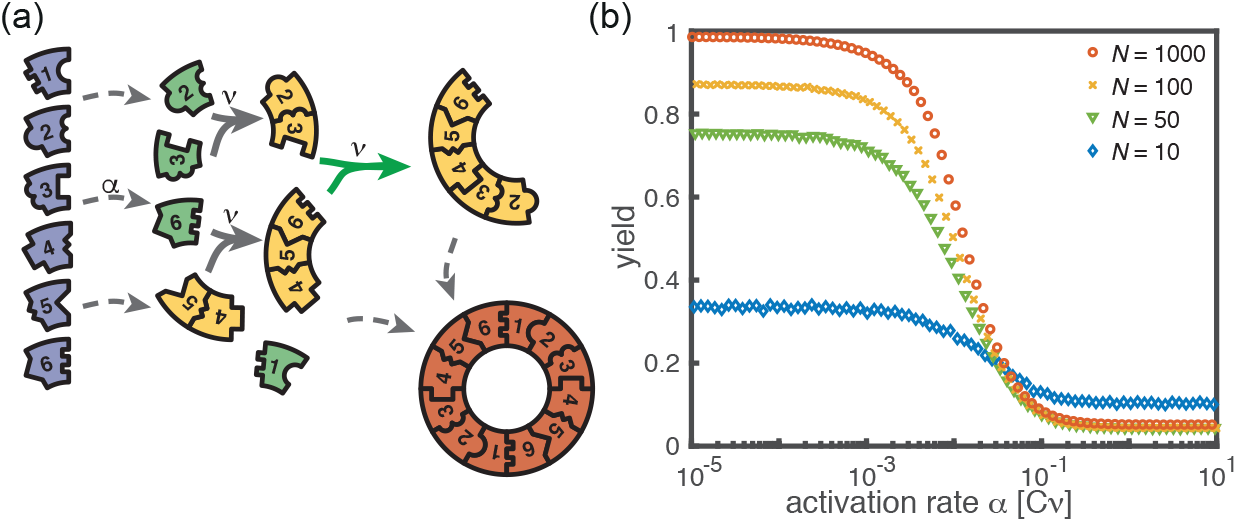
Extended model including polymer-polymer binding. (a) In the extended model, structures not only grow by monomer attachment but also by binding with another polymer (colored arrow). As before, binding happens between periodically consecutive species with rate *ν* per binding site. The reaction rate for two polymers is identical to the one for monomer-polymer binding, *ν*. Furthermore, only polymers with combined length *≤ L* can bind. All other processes and rules are the same as in the original model described in Fig. 1. (b) Dependence of the yield of the extended model on the activation rate *α* for *S* = *L* = 100, *µ* = *ν* = 1, *L*_nuc_ = 2 and different values of the number of particles per species, *N*. The qualitative behavior is the same as for the original model, in particular yield saturates (in the stochastic limit) at an imperfect value for slow activation rates. Note that polymer-polymer binding results in an increase of the minimal yield for small particle numbers.

Kinetic trapping due to structural mismatches can occur in every (partially) irreversible heterogeneous assembly process with finite-sized target structure and limited resources. From our results, we thus expect a stochastic yield catastrophe to be common to such systems. In order to test this hypothesis, we simulated a variant of our model where we assemble finite sized squares via monomer attachment from a pool of initially inactive particles. Figure 6 depicts the dependence of the yield on the activation rate for a square of size *S* = 100. Also in this case, we find that the yield saturates at an imperfect value for small activation rates. This observation supports the general validity of our findings and indicates that stochastic yield catastrophes are a general phenomenon of heterogeneous self-assembling systems that occur if particle number fluctuations are non-negligible.

**Figure 6:**
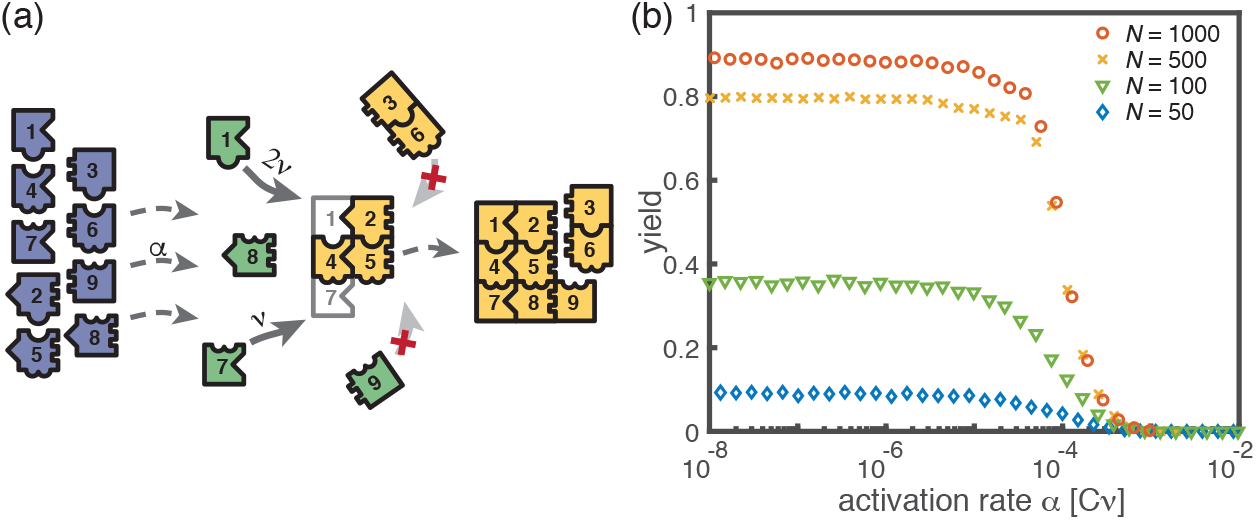
Assembly of squares of size 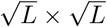 from *L* different particle species. (a) As in the ring models, there are *N* monomers of each species in the system. All particles are initially in an inactive state and are activated at the same per-capita rate *α*. Once active, species with neighboring position within the square (left/right, up/down) can bind to each other. Structures grow by attachment of single monomers until the square is complete (absorbing state). Depending on the number *b* of contacts between the monomer and the structure, the corresponding rate is *bν*. For simplicity, all polymers are stable (*L*_nuc_ = 2) and we do not consider polymer-polymer binding. (b) Dependence of the yield of the square model on the activation rate *α* for *S* = *L* = 100, *µ* = *ν* = 1 and different values of the number of particles per species, *N*. The qualitative behavior is the same as for the previous models: Whereas the yield is poor for large activation rates, it strongly increases below a threshold value and saturates (in the stochastic limit) at an imperfect value *<* 1 for small activation rates. The saturation value decreases with decreasing number of particles in the system.

## 4 Conclusions

Our results show that different ways to slow down nucleation are indeed not equivalent, and that the explicit implementation is crucial for assembly efficiency. Susceptibility to stochastic effects is highly dependent on the specific scenario. Whereas systems for which dimerization limits nucleation are robust against stochastic effects, stochastic yield catastrophes can occur in heterogeneous systems when resource supply limits nucleation. The occurrence of stochastic yield catastrophes is not captured by the deterministic rate equations, for which the qualitative behavior of both scenarios is the same. Therefore, a stochastic description of the self-assembly process, which includes fluctuations in the availability of the different species, is required. The interplay between stochastic and deterministic dynamics can lead to a plethora of interesting behaviors. For example, the combination of slow activation and slow nucleation may result in a non-monotonic dependence of the yield on the activation rate. While deterministically, yield is always improved by decreasing the activation rate, stochastic fluctuations between species strongly suppress the yield for small activation rate by effectively enhancing the nucleation speed. This observation clearly demonstrates that a *deterministically* slow nucleation speed is not sufficient in order to obtain good yield in heterogeneous self-assembly. For example, a slow activation step does not necessarily result in few nucleation events although deterministically this behavior is expected. Thus, our results indicate that the slow nucleation principle has to be interpreted in terms of the stochastic framework and have important implications for yield optimization.

We showed that demographic noise can cause stochastic yield catastrophes in heterogeneous self-assembly. However, other types of noise, such as spatiotemporal fluctuations induced by diffusion, are also expected to trigger stochastic yield catastrophes. Hence, our results have broad implications for complex biological and artificial systems, which typically exhibit various sources of noise. We characterize conditions under which stochastic yield catastrophes occur, and demonstrate how they can be mitigated. These insights could usefully inform the design of experiments to circumvent yield catastrophes: In particular, while slow provision of constituents is a feasible strategy for experiments, it is highly susceptible to stochastic effects. On the other hand, irrespective of its robustness to stochastic effects, the experimental realization of the dimerization scenario relies on cooperative or allosteric effects in binding, and may therefore require more sophisticated design of the constituents ^39, 40^. Our theoretical analysis shows that stochasticity can be alleviated either by decreasing heterogeneity (presumably lowering realizable complexity) or by increasing reversibility (potentially requiring fine-tuning of binding affinities). Alternative approaches include the promotion of specific assembly paths ^28, 41^ and the control of fluctuations ^42^. Moreover, these ideas are relevant for the understanding of intracellular self-assembly. In cells, provision of building blocks is typically a gradual process, as synthesis is either inherently slow or an explicit activation step, such as phosphorylation, is required. In addition, the constituents of the complex structures assembled in cells are usually present in small numbers and subject to diffusion. Hence, stochastic yield catastrophes would be expected to have devastating consequences for self-assembly, unless the relevant cellular processes use elaborate control mechanisms to circumvent stochastic effects. Further exploration of these control mechanisms should enhance the understanding of self-assembly processes in cells and help improve synthesis of complex nanostructures.

## 5 Methods

Here we show the derivation of Eq. 1 in the main text, giving the threshold values for the rate constants below which finite yield is obtained. The details can be found in the SI.

### Master equation and chemical rate equations

We start with the general Master equation and derive the chemical rate equations (deterministic/mean-field equations) for the heterogeneous self-assembly process. We renounce to show the full Master equation here but instead state the system that describes the evolution of the first moments. To this end, we denote the random variable that describes the number of polymers of size *ℓ* and species *s* in the system at time *t* by 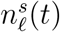 with 2 ≤ *ℓ* < *L* and 1 ≤ *s* ≤ *S*. The species of a polymer is defined by the species of the respective monomer at its left end. Furthermore, 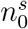 and 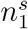 denote the number of inactive and active monomers of species *s*, respectively, and *n*_*L*_ the number of complete rings. We signify the reaction rate for binding of a monomer to a polymer of size *ℓ* by *ν*_*ℓ*_. *α* denotes the activation rate and *δ_ℓ_* the decay rate of a polymer of size *ℓ*. By 〈…〉 we indicate (ensemble) averages. The system governing the evolution of the first moments (the averages) of the 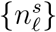 is then given by:

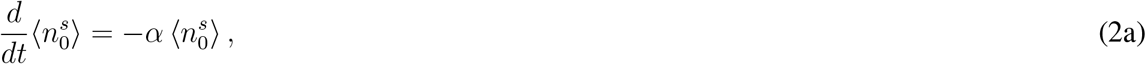

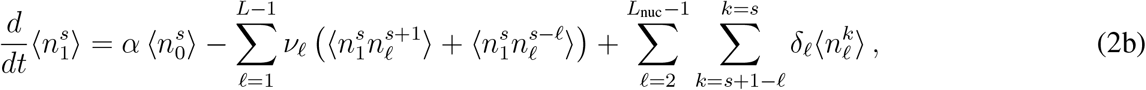

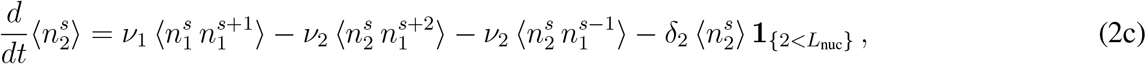

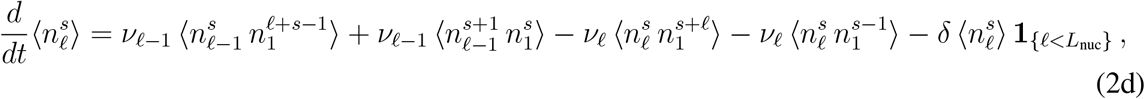

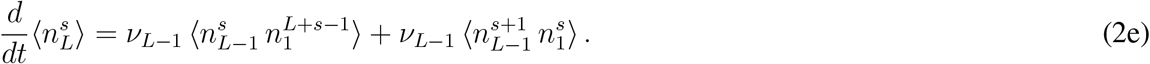

The first equation describes loss of inactive particles due to activation at rate *α*. Eq. (2b) gives the temporal change of the number of active monomers that is governed by the following processes: activation of inactive monomers at rate *α*, binding of active monomers to the left or to the right end of an existing structure of size *ℓ* at rate *ν*_*ℓ*_, and decay of below-critical polymers of size *ℓ* into monomers at rate *δ*_*ℓ*_ (disassembly).

Equations (2c) and (2d) describe the dynamics of dimers and larger polymers of size 3 *≤ ℓ < L*, respectively. The terms account for reactions of polymers with active monomers (polymerization) as well as decay in the case of below-critical polymers (disassembly). The indicator function **1**_{*x<L*__nuc}_equals 1 if the condition *x* < *L*_nuc_ is satisfied and 0 otherwise. Note that a polymer of size *ℓ* ≥ 3 can grow by attaching a monomer to its left or to its right end whereas the formation of a dimer of a specific species is only possible via one reaction pathway (dimerization reaction). Finally, polymers of length *L* – the complete ring structures – form an absorbing state and, therefore, include only the respective gain terms (cf. Eq 2e).

We simulated the Master equation underlying Eq. (2) stochastically using Gillespie’s algorithm. For the following deterministic analysis, we neglect correlations between particle numbers 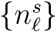, which is valid assumption for large particle numbers. Then the two-point correlator can be approximated as the product of the corresponding mean values (mean-field approximation)

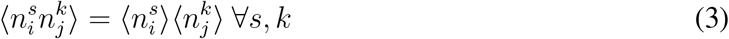

Furthermore, for the expectation values it must hold

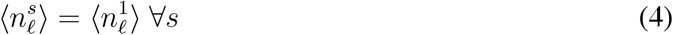

because all species have equivalent properties (there is no distinct species) and hence the system is invariant under relabelling of the upper index. By

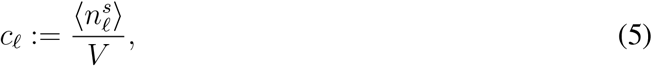

we denote the concentration of any monomer or polymer species of size *ℓ*, where *V* is the reaction volume. Due to the symmetry formulated in Eq. (4), the heterogeneous assembly process decouples into a set of *S* identical and independent homogeneous assembly processes in the deterministic limit. The corresponding homogeneous system then is described by the following set of equations that is obtained by applying (3), (4) and (5) to (2)

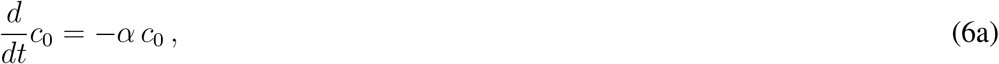

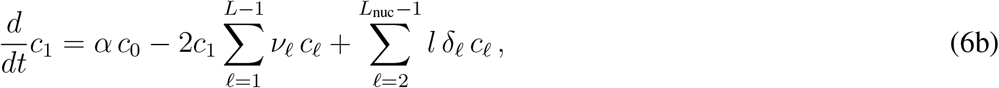

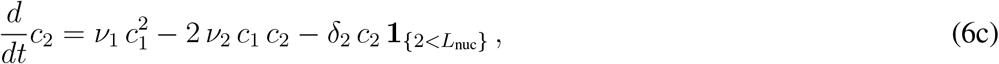

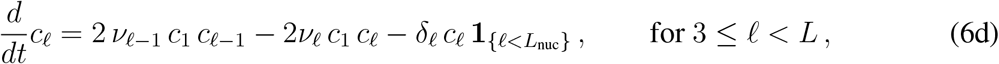

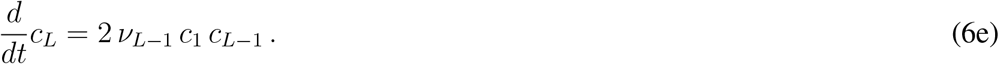

The rate constants *ν*_*ℓ*_ in Eq. (6) and (2) differ by a factor of *V*. For convenience, we use however the same symbol in both cases. The rate constants *ν*_*ℓ*_ in Eq. (6) can be interpreted in the usual units 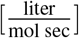. Due to the symmetry, the yield, which is given by the quotient of the number of completely assembled rings and the maximum number of complete rings, becomes independent of the number of species *S*

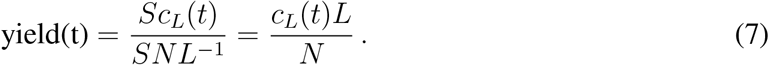

Hence, it is enough to study the dynamics of the homogeneous system, Eq. (6), to identify the condition under which non zero yield is obtained.

### Effective description by an advection-diffusion equation

The dynamical properties of the evolution of the polymer-size distribution become evident if the set of ODEs (6) is rewritten as a partial differential equation. This approach was previously described in the context of virus capsid assembly^6, 33^.

For simplicity, we restrict ourselves to the case *L*_nuc_ = 2 and let *ν*_1_ = *µ* and *ν*_*ℓ*≥2_ = *ν*. Then, for the polymers with *ℓ* > 2 we have

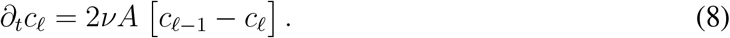

As a next step, we approximate the index *ℓ* ∈ {2, 3, …, *L*} indicating the length of the polymer as a continuous variable *x* ∈ [2, *L*] and define *c*(*x* = *ℓ*):= *c*_*ℓ*_. The concentration of active monomers is denoted by *A*. Formally expanding the right-hand side of Eq. (8) in a Taylor series up to second order

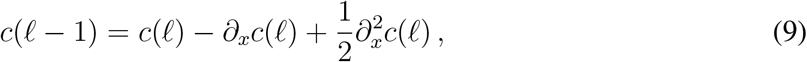

one arrives at the advection-diffusion equation with both advection and diffusion coefficients depending on the concentration of active monomers *A*(*t*)

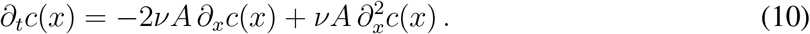

Equation (10) can be written in the form of a continuity equation *∂*_*t*_*c*(*x*) = − *∂*_*x*_*J*(*x*) with flux *J* = 2*νA c* − *νA ∂*_*x*_*c*. The flux at the left boundary *x* = 2 equals the influx of polymers due to dimerization of free monomers *J* (2*, t*) = *µA*^2^. This enforces a Robin boundary condition at *x* = 2

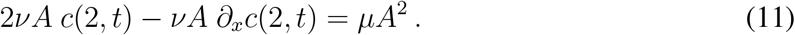

At *x* = *L* we set an absorbing boundary *c*(*L, t*) = 0 so that completed structures are removed from the system. The time evolution of the concentration of active monomers is given by

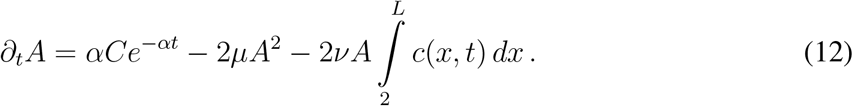

The terms on the right-hand side account for activation of inactive particles, dimerization, and binding of active particles to polymers (polymerization).

Qualitatively, Eq. (10) describes a profile that emerges at *x* = 2 from the boundary condition Eq. (11) moves to the right with time-dependent velocity 2*νA*(*t*) due to the advection term, and broadens with a time-dependent diffusion coefficient *νA*(*t*). In the SI we show how the full solution of Eqs. (10) and (11) can be found assuming knowledge of *A*(*t*). Here, we focus only on the derivation of the threshold activation rate and threshold dimerization rate that mark the onset of non-zero yield.

Yield production starts as soon as the density wave reaches the absorbing boundary at *x* = *L*. Therefore, finite yield is obtained if the sum of the advectively travelled distance *d*_adv_ and the diffusively travelled distance *d*_diff_ exceeds the system size *L* − 2

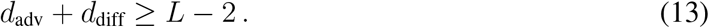

According to Eq. (10), 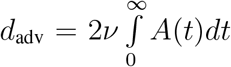 and 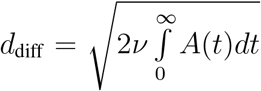, giving as condition for the onset of finite yield

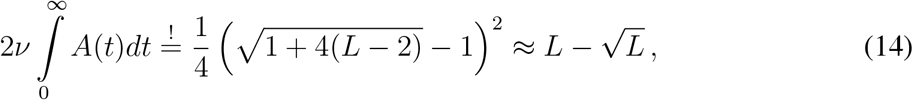

where the last approximation is valid for large *L*.

In order to obtain 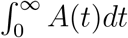 we derive an effective two-component system that governs the evolution of *A*(*t*). To this end, we denote the total number of polymers in Eq. (12) by 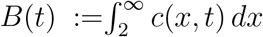 (as long as yield is zero the upper boundary is irrelevant and we can consider *L* = ∞). Eq. (12) then reads

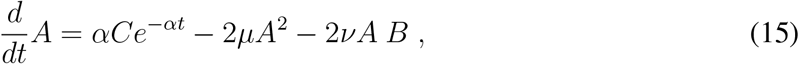

and the dynamics of *B* is determined from the boundary condition, Eq. (11)

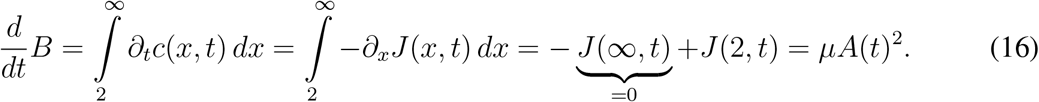

Measuring *A* and *B* in units of the initial monomer concentration *C* and time in units of (*νC*)^*−*1^ the equations are rewritten in dimensionless units as

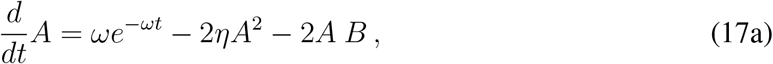

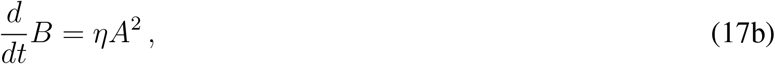

where 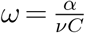 and 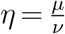. Eq. (17) describes a closed two-component system for the concentration of active monomers *A* and the total concentration of polymers *B*. It describes the dynamics exactly as long as yield is zero. In order to evaluate the condition (14) we need to determine the integral over *A*(*t*) as a function of *ω* and *η*

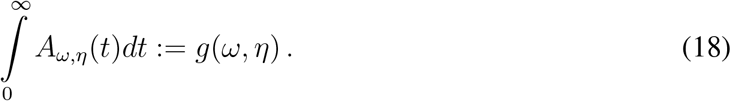

To that end, we proceed by looking at both scenarios separately. The numerical analysis, confirming our analytic results, is given in the SI.

### Dimerization scenario

The activation rate in the dimerization scenario is *α* → *∞*, and instead of the term *ωe*^−*ωt*^ in d*A/*d*t*, we set the initial condition *A*(0) = 1 (and *B*(0) = 0). Furthermore, *η* = *µ/ν ≪* 1 and we can neglect the term proportional to *η* in d*A/*d*t*. As a result,

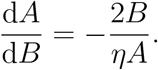

Solving this equation for *A* as a function of *B* using the initial condition *A*(*B* = 0) = 1, the totally travelled distance of the wave is determined to be

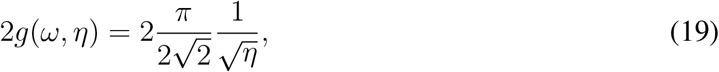

where for the evaluation of the integral we used the substitution *ηA*^2^d*t* = d*B*.

### Activation scenario

In the activation scenario, yield sets in only if the activation rate and thus the effective nucleation rate is slow. As a result, in addition to *ω* ≪ 1, we can again neglect the term proportional to *η* in d*A/*d*t*. This time, however, we have to keep the term *ωe*^−*ωt*^. As a next step, we assume that d*A/*d*t* is much smaller than the remaining terms on the right-hand side, *ωe*^−*ωt*^ and *−*2*AB*. This assumption might seem crude at first sight but is justified *a posteriori* by the solution of the equation (see SI). Hence, we get the algebraic equation *A*(*t*) = *ωe*^−*ωt*^/(2*B*(*t*)). Using it to solve d*B/*d*t* = *ηA*^2^ for *B*, and then to determine *A*, the totally travelled distance of the wave is deduced as

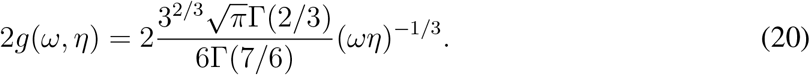

Taken together, we therefore obtain two conditions out of which one must be fulfilled in order to obtain finite yield

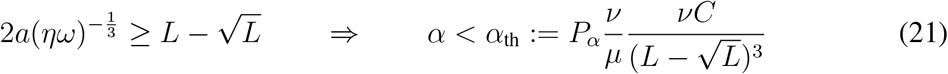

or

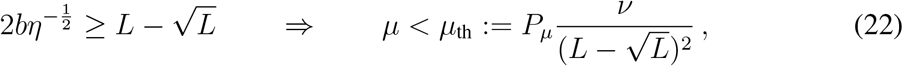

where *a* and *b* are numerical factors, and *P*_*α*_ = 8*a*^3^ ≈ 5.77 and *P*_*µ*_ = 4*b*^2^ ≈ 4.93. This verifies Eq. (1) in the main text.

## Acknowledgements

We thank Nigel Goldenfeld for a stimulating discussion, and Raphaela Geßele and Laeschkir Hassan for helpful feedback on the manuscript. This research was supported by the German Excellence Initiative via the program ‘NanoSystems Initiative Munich’(NIM). F.M.G. and I.R.G. are supported by a DFG fellowship through the Graduate School of Quantitative Biosciences Munich (QBM). We also gratefully acknowledge financial support by the DFG Research Training Group GRK2062 (Molecular Principles of Synthetic Biology). Finally, E.F. thanks the Aspen Center for Physics, which is supported by National Science Foundation grant PHY-1607611, for their hospitality and inspiring discussions with colleagues.

