## Supplemental Information for "Stochastic Yield Catastrophes and Robustness in Self-Assembly"

### Supplementary Information

#### S1 Chemical reaction equations and the equivalence of models with different numbers of species

In this section we derive the chemical rate equations (deterministic equations) for the self-assembly process as described in the main text. Furthermore, we show that for general $S$ in the deterministic limit the model is equivalent to a set of $S$ independent assembly processes with only one species.

**Homogeneous structures** First, we consider the homogeneous model ($S=1$). By $c_\ell(t)$ we denote the concentration of complexes of length $\ell$ ($\ell \geq 2$) at time $t$, $c_1(t)$ is the concentration of active monomers and $c_0(t)$ the concentration of inactive monomers at time $t$. In the following we will usually skip the time argument for better readability. We denote the reaction rate for binding of a monomer to a polymer of size $\ell$ by $\nu_\ell$. The model from the main text is recovered by setting $\nu_\ell := \mu_\ell$ if $\ell < L_{\text{nuc}}$, and $\nu_\ell := \nu$ otherwise. The ensuing set of ordinary differential equations then reads:

$$\frac{d}{dt}c_0 = -\alpha\, c_0\,, \tag{S1a}$$

$$\frac{d}{dt}c_1 = \alpha\, c_0 - 2c_1 \sum_{\ell=1}^{L-1} \nu_\ell\, c_\ell + \sum_{\ell=2}^{L_{\text{nuc}}-1} l\, \delta_\ell\, c_\ell\,, \tag{S1b}$$

$$\frac{d}{dt}c_2 = \nu_1\, c_1^2 - 2\,\nu_2\, c_1\, c_2 - \delta_2\, c_2\, \mathbf{1}_{\{2<L_{\text{nuc}}\}}\,, \tag{S1c}$$

$$\frac{d}{dt}c_\ell = 2\,\nu_{\ell-1}\, c_1\, c_{\ell-1} - 2\nu_\ell\, c_1\, c_\ell - \delta_\ell\, c_\ell\, \mathbf{1}_{\{\ell<L_{\text{nuc}}\}}\,, \qquad \text{for } 3 \leq \ell < L\,, \tag{S1d}$$

$$\frac{d}{dt}c_L = 2\,\nu_{L-1}\, c_1\, c_{L-1}\,. \tag{S1e}$$

The indicator function $\mathbf{1}_{\{x<L_{\text{nuc}}\}}$ equals 1 if the condition $x < L_{\text{nuc}}$ is satisfied and 0 otherwise. The first equation describes loss of inactive particles due to activation at rate $\alpha$. The equation is uncoupled from the remainder of the equations and is solved by $c_0(t) = Ce^{-\alpha t}$, with $C$ denoting the initial concentration of inactive monomers. The temporal change of the active monomers is governed by the following processes (Eq. (S1b)): activation of inactive monomers at rate $\alpha$, binding of active monomers to existing structures at rate $\nu_\ell$ (polymerization), and decay of below-critical polymers into monomers at rate $\delta_\ell$ (disassembly). All binding rates appear with a factor of 2 because a monomer can attach to a polymer on its left or on its right end.

Note that there is a subtlety with the dimerization term "$2\,\nu_1\, c_1^2$": the dimerization term as well bears a factor of 2 because two identical monomers $A$ and $B$ can form a dimer in two possible ways, either as $AB$ or $BA$. Additionally, there is a stoichiometric factor of 2 for this reaction.

However, one factor of 2 is cancelled again because, assuming there are $n$ monomers, the number of ordered pairs of monomers that describe possible reaction partners is $\frac{1}{2}n(n-1) \approx n^2/2$ (if $n$ is large) rather than $n^2$ (the number of reaction partners when two different species react). This leaves us with a single factor of 2 like for all the other binding reactions.

Equations (S1c) and (S1d) describe the dynamics of dimers and larger polymers of size $3 \leq \ell < L$, respectively. The terms account for reactions of polymers with active monomers (polymerization) as well as decay in the case of below-critical polymers (disassembly). The dimerization term in the equation for $\partial_t c_2$ lacks the factor of 2 because the stoichiometric factor is missing as compared with the dimerization term in the line above. Finally, polymers of length $L$ – the complete ring structures – form an absorbing state and therefore only include a reactive gain term (Eq. (S1e)).

**Heterogeneous structures** Next we consider systems with more than one particle species ($S > 1$). The heterogeneous system can be described by dynamical equations equivalent to the homogeneous system. We show this starting from a full description that distinguishes both monomers and polymers into a set of different species $1, \ldots, S$. In order to formulate the dynamic equations and to see the equivalence to a one-species model, we distinguish both monomers and polymers into a set of different species $1, \ldots, S$. The species of a polymer is defined by the species of the respective monomer at its left end. As polymers assemble in consecutive order of species, a polymer is uniquely determined by its length and species (i.e. species of leftmost monomer). In that sense, $c_\ell^s$ with $0 \leq \ell < L$ and $1 \leq s \leq S$ denotes the concentration of a polymer of length $\ell$ and species $s$ ($c_0^s$ and $c_1^s$ again denote inactive and active monomers of species $s$, respectively). For example, $c_4^5$ denotes the concentration of polymers $[5678]$ if $S \geq 8$, or of polymers $[5612]$ if $S = 6$. Upper indices are always assumed to be taken modulo $S$ whenever they lie outside the range $[1, S]$. Therefore, the dynamics of the concentrations $c_\ell^s$ with $3 \leq \ell < L$ is given by

$$\frac{d}{dt} c_\ell^s = \nu_{\ell-1}\, c_{\ell-1}^s\, c_1^{\ell+s-1} + \nu_{\ell-1}\, c_{\ell-1}^{s+1}\, c_1^s - \nu_\ell\, c_\ell^s\, c_1^{s+\ell} - \nu_\ell\, c_\ell^s\, c_1^{s-1} - \delta\, c_\ell^s\, \mathbf{1}_{\{\ell < L_{\text{nuc}}\}} \,. \tag{S2}$$

The terms on the right-hand side account for the influx due to binding of the respective polymers of length $\ell - 1$ with a monomer either on the right or on the left (first and second term), and for the outflux due to reactions of a polymer of length $\ell$ and species $s$ (third and fourth term), as well as for decay into monomers for $\ell < L_{\text{nuc}}$ (last term). For the dynamics of the dimers, however, there is only one gain term arising from dimerization:

$$\frac{d}{dt} c_2^s = \nu_1\, c_1^s\, c_1^{s+1} - \nu_2\, c_2^s\, c_1^{s+2} - \nu_2\, c_2^s\, c_1^{s-1} - \delta_2\, c_2^s\, \mathbf{1}_{\{2 < L_{\text{nuc}}\}} \,. \tag{S3}$$

Equivalently, for the active monomers we find:

$$\frac{d}{dt}c_1^s = \alpha\, C e^{-\alpha t} - c_1^s \sum_{\ell=1}^{L-1} \nu_\ell \left(c_\ell^{s+1} + c_\ell^{s-\ell}\right) + \sum_{\ell=2}^{L_{\text{nuc}}-1} \sum_{k=s+1-\ell}^{k=s} \delta_\ell c_\ell^k \,. \tag{S4}$$

Now we exploit the symmetry of the system with respect to the species index, that is, the upper index in $\{c_\ell^s\}$: Since all species in the system are equivalent, the dynamic equations are invariant under relabelling of the upper indices. Consequently, it must hold that:

$$c_\ell^s(t) = c_\ell^k(t), \qquad \text{for any } s, k \leq S \text{ at any time } t. \tag{S5}$$

In other words, the upper index is irrelevant and can also be discarded. The variable $c_\ell$ then denotes the concentration of *any* one polymer species of length $\ell$. Taking advantage of this symmetry for the equations of the heterogeneous system, (Eq. (S2), Eq. (S3) and Eq. (S4)), and collecting equal terms leads to a set of equations fully identical to those for the homogeneous system (Eq. (S1)). We show the equivalence to the homogeneous model exemplarily for the dynamics of the polymers with size $\ell \geq 3$ in Eq. (S2). Applying $c_\ell^s(t) = c_\ell(t)$ to Eq. (S2) yields for the dynamics of the concentration of an arbitrary polymer species of size $\ell$:

$$\begin{aligned}\frac{d}{dt}c_\ell &= \nu_{\ell-1}\, c_{\ell-1}\, c_1 + \nu_{\ell-1}\, c_{\ell-1}\, c_1 - \nu_\ell\, c_\ell\, c_1 - \nu_\ell\, c_\ell\, c_1 - \delta\, c_\ell\, \mathbf{1}_{\{\ell < L_{\text{nuc}}\}}\,. \\ &= 2\nu_{\ell-1}\, c_{\ell-1}\, c_1 - 2\nu_\ell\, c_\ell\, c_1 - \delta\, c_\ell\, \mathbf{1}_{\{\ell < L_{\text{nuc}}\}},\end{aligned}$$

which is identical to the respective dynamic equation (S1d) for the homogeneous model. The other equations for the heterogeneous system reduce to those for the homogeneous system in an analogous manner.

Summarizing, we have shown that the (deterministic) heterogeneous assembly process decouples into a set of $S$ identical and independent homogeneous processes. In particular, yield, which is given by the quotient of the number of completely assembled rings and the maximal possible number of complete rings, becomes independent of $S$:

$$\text{yield(t)} = \frac{S c_L(t)}{S N L^{-1}} = \frac{c_L(t) L}{N}\,. \tag{S6}$$

#### S2 Effective description of the evolution of the polymer size distribution as an advection-diffusion equation

The dynamical properties of the evolution of the polymer size distribution become evident if the set of ODEs (S1) is rewritten as a partial differential equation. This approach was previously described in the context of virus capsid assembly[6,33] but we will restate the essential steps here for the convenience of the reader. To this end we interpret the length index of the polymer $\ell \in \{2, 3, \dots, L\}$ as a continuous variable that we rename $x \in [2, L]$. With such a continuous description in view we write $c(x = \ell) := c_\ell$ to denote the concentration of polymers of size $\ell$.

Since the active monomers play a special role, we denote their concentration in the following by $A$. For simplicity we restrict our discussion to the case $L_{\text{nuc}} = 2$ and let $\nu_1 = \mu$ and $\nu_{\ell \geq 2} = \nu$. Generalizations to $L_{\text{nuc}} > 2$ can be done in a similar way. Then, for the polymers with $\ell \geq 3$ we have:

$$\partial_t c(\ell) = 2\nu A \left[ c(\ell - 1) - c(\ell) \right] . \tag{S7}$$

Formally, expanding the right-hand side in a Taylor series up to second order

$$c(\ell - 1) = c(\ell) - \partial_x c(\ell) + \frac{1}{2} \partial_x^2 c(\ell) , \tag{S8}$$

we arrive at an advection-diffusion equation with both advection and diffusion coefficients depending on the concentration of active monomers $A(t)$,

$$\partial_t c(x) = -2\nu A \, \partial_x c(x) + \nu A \, \partial_x^2 c(x) . \tag{S9}$$

Equation (S9) can be written in the form of a continuity equation $\partial_t c(x) = -\partial_x J(x)$ with flux $J = 2\nu A \; c - \nu A \; \partial_x c$. The flux at the left boundary, $x = 2$, equals the influx of polymers due to dimerization of free monomers, $J(2, t) = \mu A^2$. This enforces a Robin boundary condition at $x = 2$,

$$2\nu A \; c(2, t) - \nu A \; \partial_x c(2, t) = \mu A^2 . \tag{S10}$$

At $x = L$, we have an absorbing boundary $c(L, t) = 0$ so that completed structures are removed from the system. Furthermore, the time evolution of the concentration of active particles is given by

$$\partial_t A = \alpha C e^{-\alpha t} - 2\mu A^2 - 2\nu A \int_2^L c(x, t) \, dx . \tag{S11}$$

The terms on the right-hand side account for activation of inactive particles, dimerization, and binding of active particles to polymers (polymerization).

Qualitatively, Eq. (S9) describes a profile that emerges at $x = 2$ from the boundary condition, Eq. (S10), moves to the right with time dependent velocity $2\nu A(t)$ due to the advection term, and broadens with a time-dependent diffusion coefficient $\nu A(t)$. The concentration of active particles $A$ determines both the influx of dimers at $x = 2$, as well as the speed and diffusion of the wave profile.

Next, we derive an expression that solves Eq. (S9), assuming that we know $A(t)$. We start by solving Eq. (S9) at the left boundary $c(2,t)$, and then translate the resulting expression to obtain a solution for $c(x,t)$. To obtain $c(2,t)$ in dependence of $a(t)$ we can solve $\frac{d}{dt}c(2,t) = \mu A^2 - \nu A c(2,t)$ (see Eq. (S1c)) by 'variation of the constants' as

$$c(2,t) = \int_0^t \mu A(\tilde{t})^2 \exp\left[-\int_{\tilde{t}}^t \nu A(t')dt'\right] d\tilde{t}\ . \tag{S12}$$

With help of this expression we find $c(x,t)$: Given $c(2,t)$, the advective part of Eq. (S9),

$$\partial_t \tilde{c}(x) = -2\nu A\, \partial_x \tilde{c}(x)\ . \tag{S13}$$

is solved by

$$c_{\text{advec}}(x,t) = c(2, \tau(x,t))\ . \tag{S14}$$

Here, $\tau(x,t)$ denotes the time that a particle at position $x$ and time $t$ was at $x = 2$. In other words, a particle at time $t$ and position $x$ has entered the system at $x = 2$ at time $\tau(x,t)$. This ansatz solves the PDE (Eq. (S13)) if and only if $\tau(x,t)$ satisfies

$$\tau(x,t) = \tilde{A}^{-1}\left(\tilde{A}(t) - \frac{x-2}{2\nu}\right) \tag{S15}$$

with $\tilde{A}$ being an arbitrary integral of $A$ such that $\partial_t \tilde{A}(t) = A(t)$ and $\tilde{A}^{-1}$ denoting its inverse. More easily, we find this form of $\tau$ by requiring that the integral over the velocity from time $\tau$ to $t$ equals the travelled distance $x - 2$:

$$\int_\tau^t 2\nu\, A(t')dt' = x - 2\ . \tag{S16}$$

To include the diffusive contribution in Eq. (S13), we use the diffusion kernel,

$$k(x,y,t) = \left(4\pi \int_{\tau(y,t)}^t D(t)\right)^{-1/2} \exp\left(\frac{-x^2}{4\int_{\tau(y,t)}^t D(t)}\right), \tag{S17}$$

with the time dependent diffusion constant $D(t) = \nu A(t)$. The kernel $k(x,y,t)$ accounts for the mass that has been diffusively transported from $y$ a distance of $x$. Because the mass has entered the system at $x = 2$ at time $\tau(y,t)$, it diffused for the time $t - \tau(y,t)$. The complete expression for $c(x,t)$ is then obtained as the convolution of $c_{\text{advec}}(x,t)$ (Eq. (S14)), that is obtained from Eq. (S12) and Eq. (S15), and the diffusion kernel $k(x,y,t)$ (Eq. (S17)):

$$c(x,t) = \int c_{\text{advec}}(s,t) k(x-s,s,t)ds = \int c(2,\tau(s,t))k(x-s,s,t)ds\ . \tag{S18}$$

Interpreting the terms in the equations and the general form of the solution, we are able to understand the qualitative behavior of the system. If both the activation and the dimerization rate are large, the system produces zero yield: both advection and diffusion are driven by the concentration of active monomers $A$. If activation is fast, the concentration of active monomers $A$ will become large initially since activation is faster than the reaction dynamics. Consequently, provided $\mu \sim \nu$, dimerization dominates over binding because it depends quadratically on $A$, see Eq. (S11). The reservoir of free particles then depletes quickly and cannot sustain the motion of the wave for long enough to reach the absorbing boundary, resulting in a very low yield. Only if either the activation rate is low enough or if $\mu \ll \nu$, the motion of the wave can be sustained until it reaches the absorbing boundary.

#### S3 Threshold values for the activation and dimerization rate

Based on the analysis from the previous section, we will now determine the threshold activation rate and threshold dimerization rate which mark the onset of non-zero yield. Yield production starts as soon as the density wave reaches the absorbing boundary at $x = L$. Therefore, finite yield is obtained if and only if the sum of the advectively travelled distance $d_{\text{adv}}$ and the diffusively travelled distance $d_{\text{diff}}$ exceeds the system size $L - 2$:

$$d_{\text{adv}} + d_{\text{diff}} \geq L - 2 \,. \tag{S19}$$

The condition for the onset of non-zero yield is obtained by assuming equality in this relation. The advectively travelled distance is obtained from Eq. (S16) by setting the borders of the integral over the velocity to $\tau = 0$ and $t = \infty$:

$$d_{\text{adv}} = \int_0^\infty 2\nu A(t')dt'. \tag{S20}$$

The diffusively travelled distance is approximately given by the standard deviation of the Gaussian diffusion kernel, Eq. (S17), again with $\tau = 0$ and $t = \infty$,

$$d_{\text{diff}} = \sqrt{2\nu \int_0^\infty A(t)dt}. \tag{S21}$$

Taken together, we obtain a condition for the onset of finite yield:

$$2\nu \int_0^\infty A(t)dt + \sqrt{2\nu \int_0^\infty A(t)dt} = L - 2 \,. \tag{S22}$$

Substituting $y = \sqrt{2\nu \int A}$ and requiring that $y$ is positive, we can solve the quadratic equation and find that Eq. (S22) is equivalent to

$$2\nu \int_0^\infty A(t)dt = y^2 = \frac{1}{4}\left(\sqrt{1+4(L-2)} - 1\right)^2 \approx L - \sqrt{L}\,, \tag{S23}$$

where the last approximation is valid for large $L$.

We determine the threshold values for the activation rate $\alpha$ and the dimerization rate $\mu$ by finding solutions of the dynamical equation for the active particles $A(t)$, Eq. (S11), such that the condition, Eq. (S23), is fulfilled. Thus, we start by deriving the dependence of $\int_0^\infty A(t)dt$ on $\alpha$ and $\mu$.

The concentration $c(x,t)$ appears in Eq. (S11) only in terms of an integral $\int_2^L c(x,t)\,dx$, counting the total number of polymers in the system. As long as yield is zero there is no outflux of polymers at the absorbing boundary $x = L$ and the total number of polymers in the system only increases due to the influx at the left boundary $x = 2$. As long as yield is zero we can therefore equivalently consider the limit $L \to \infty$. We denote the total number of polymers in Eq. (S11) by $B(t) := \int c(x,t)\,dx$ for which the dynamics is determined from the boundary condition, Eq. (S10):

$$\frac{d}{dt}B = \int_2^\infty \partial_t c(x,t)\,dx = \int_2^\infty -\partial_x J(x,t)\,dx = -\underbrace{J(\infty,t)}_{=0} + J(2,t) = \mu A(t)^2. \tag{S24}$$

Hence, as long as yield is zero, the total number of polymers increases with the rate of the dimerization events. The system then simplifies to a set of two coupled ordinary differential equations for $A$ and $B$:

$$\frac{d}{dt}A = \alpha C e^{-\alpha t} - 2\mu A^2 - 2\nu A\,B\,, \tag{S25a}$$

$$\frac{d}{dt}B = \mu A^2\,. \tag{S25b}$$

The dynamics of $A$ and $B$ is equivalent to a two-state activator-inhibitor system, where $A$ dimerizes into $B$ at rate $\mu$, and $B$ degrades (inhibits) $A$ at rate $2\nu$. Note that Eq. (S25a) describes the exact dynamics of the active monomers $A$ and total number of polymers $B$ in the deterministic system as long as yield is zero. The system has therefore been greatly reduced from originally $S\,N$ coupled ODEs to now only 2 coupled ODEs.

For the further analysis it is useful to non-dimensionalize Eq. (S25a) by measuring $A$ and $B$

in units of the initial concentration of inactive monomers $C$ and time in units of $(\nu C)^{-1}$:

$$\frac{d}{dt}A = \omega e^{-\omega t} - 2\eta A^2 - 2A\,B\,, \tag{S26a}$$

$$\frac{d}{dt}B = \eta A^2\,, \tag{S26b}$$

with the remaining dimensionless parameters $\omega = \frac{\alpha}{\nu C}$ and $\eta = \frac{\mu}{\nu}$. We are interested in the integral over $A(t)$ as a function of $\omega$ and $\eta$,

$$\int_0^\infty A_{\omega,\eta}(t)dt := g(\omega, \eta)\,, \tag{S27}$$

which relates to the totally travelled distance of the wave. Note that, in case of zero yield, $2g(\omega, \eta)$ is the total advectively travelled distance of the wave (cf. Eq. (S20)) and the square of the diffusively travelled distance (cf. Eq. (S21)).

**Analysis of the dimerization scenario** The dimerization scenario is characterized by fast activation $\alpha \gg C\nu$ and slow dimerization $\mu \ll \nu$. For the dimensionless parameters these assumptions translate to $\eta \ll 1$ and $\eta \ll \omega$. Because for small $\eta \ll 1$ nucleation is much slower than growth we neglect the dimerization term in Eq. (S26a) against the growth term. Furthermore, because $\eta \ll \omega$ activation happens on a fast time scale compared with nucleation and we may therefore integrate out the fast time scale assuming that all particles are activated instantaneously at the beginning. The system Eq. (S26) then reduces to

$$\frac{d}{dt}A = -2A\,B\,, \tag{S28a}$$

$$\frac{d}{dt}B = \eta A^2\,, \tag{S28b}$$

with the initial condition $A(0) = 1$ and $B(0) = 0$. We divide the first equation by the second one (formally applying the chain rule and the inverse function theorem) to obtain a single equation for the dynamics of $A(B)$:

$$\frac{dA}{dB} = -\frac{2}{\eta}\frac{B}{A}\,, \tag{S29}$$

where $A(B{=}0) = 1$. This first order ODE can be solved by separation of variables and subsequent integration, yielding

$$A(B) = \sqrt{1 - \frac{2}{\eta}B^2}\,. \tag{S30}$$

Because the number of active monomers $A(t)$ must vanish for $t \to \infty$, the final value of $B$ is

$$B_\infty := B(t{=}\infty) = \sqrt{\frac{\eta}{2}}\,. \tag{S31}$$

Thereby, we calculate the function $g(\eta)$ via variable substitution $dt = \frac{dB}{\eta A^2}$:

$$g(\eta) = \int_0^\infty A(t)dt = \int_0^{B_\infty} A(B)\frac{dB}{\eta A(B)^2} = \frac{1}{\eta}\int_0^{B_\infty} \frac{dB}{\sqrt{1-\frac{2}{\eta}B^2}} = \frac{\pi}{2\sqrt{2}}\,\eta^{-\frac{1}{2}}\,. \tag{S32}$$

So, the dependence of the travelled distance of the wave on $\eta$ obeys a power law with exponent $-\frac{1}{2}$, confirming the previous result[33]. For the coefficient we find $\frac{\pi}{2\sqrt{2}} \approx 1.1107$.

Additionally, we can determine the time dependent solutions $A(t)$ and $B(t)$. Using the solution for $A(B)$ from Eq. (S30) in Eq. (S28b) we obtain $B(t)$ as

$$B(t) = \sqrt{\frac{\eta}{2}}\tanh\left(\sqrt{2\eta}t\right)\,. \tag{S33}$$

We use this expression for $B(t)$ in Eq. (S28a) to obtain $A(t)$. The resulting ODEs can again be solved by separation of variables as

$$A(t) = \frac{1}{\cosh\left(\sqrt{2\eta}t\right)}. \tag{S34}$$

**Analysis of the activation scenario** In the activation scenario, $\alpha \ll C\nu$, such that $\omega \ll 1$ and $\omega \ll \eta$. As we know already that decreasing $\omega$ will slow down nucleation relative to growth we can again neglect the dimerization term in Eq. (S26a). In contrast to the dimerization scenario, however, we have to keep the activation term. Transforming time via $\tau := 1 - e^{-\omega t}$ such that $\tau \in [0,1]$ and writing $a(\tau) = a(1-e^{-\omega t}) := A(t)$ and $b(\tau) = b(1-e^{-\omega t}) := B(t)$ the system in Eq. (S26) becomes:

$$\frac{d}{d\tau}a = 1 - \frac{2}{\omega(1-\tau)}ab\,, \tag{S35a}$$

$$\frac{d}{d\tau}b = \frac{\eta}{\omega(1-\tau)}a^2\,, \tag{S35b}$$

with the initial condition $a(0) = b(0) = 0$. The function $g(\omega,\eta)$ transforms as

$$g(\omega,\eta) = \int_0^\infty A(t)dt = \frac{1}{\omega}\int_0^1 \frac{a(\tau)}{1-\tau}d\tau. \tag{S36}$$

In the following we derive the asymptotic solution for $a(\tau)$ in the limit of small $\omega$ in order to evaluate the integral in Eq. (S36). In the limit $\tau \to 1$ ($\Leftrightarrow t \to \infty$) both $a(\tau)$ and $\frac{d}{d\tau}a(\tau)$ will become small whereas $b(\tau)$ increases monotonically. The reaction term in Eq. (S35a) is furthermore weighted by a factor $\frac{1}{\omega}$ which will become large if $\omega \ll 1$. We therefore postulate that for sufficiently large $\tau$ the derivative $\frac{d}{d\tau}a(\tau)$ is much smaller than the two terms on the right-hand side of Eq. (S35a) and hence negligible. This assumption has to be justified *a posteriori* with the obtained solution. Neglecting the derivative term $\frac{d}{d\tau}a$ in (S35a) reduces the equation to an algebraic equation and we find

$$a = \frac{\omega(1-\tau)}{2b}. \tag{S37}$$

Using this result in Eq. (S35b) we can solve for $b$ by separation of variables and subsequent integration:

$$b(\tau) = (\omega\eta)^{\frac{1}{3}} \cdot \left(\frac{3}{4}\tau - \frac{3}{8}\tau^2\right)^{\frac{1}{3}} . \tag{S38}$$

From Eq. (S37) we immediately obtain $a(\tau)$:

$$a(\tau) = \frac{\omega^{\frac{2}{3}}}{\eta^{\frac{1}{3}}} \cdot \frac{1-\tau}{(6\tau - 3\tau^2)^{\frac{1}{3}}} := \frac{\omega^{\frac{2}{3}}}{\eta^{\frac{1}{3}}} h(\tau) , \tag{S39}$$

where by $h(\tau)$ we denote the part of the solution that depends only on $\tau$. Hence, we find that $a$ and hence also $\frac{d}{d\tau}a$ scale like $\sim \omega^{\frac{2}{3}}$, and will thus become small if $\omega \ll 1$ and $\tau$ is large enough. Therefore the solution is consistent[1] and justifies the approximation in which we neglected the derivative term in the limit of small $\omega$ and sufficiently large $\tau$.

[1]Consistency of the solution with the approximation is a sufficient criterion for the validity of the approximation: We can solve the system for $A$ and $B$ in Eq. (S35) iteratively by defining

$$\frac{d}{d\tau}a_{i-1} = 1 - \frac{2}{\omega(1-\tau)}a_i b_i ,$$
$$\frac{d}{d\tau}b_i = \frac{\eta}{\omega(1-\tau)}a_i^2 .$$

Assuming that for $i \to \infty$, $a_i$ and $b_i$ converge to the correct solutions $a(\tau)$ and $b(\tau)$ when starting with $a_0 = 0$, we obtain $a_1$ and $b_1$ as given by Eq. (S39) and Eq. (S38) and can iteratively refine the approximation. The next iteration step then reads: $\frac{d}{d\tau}a_1 = 1 - \frac{2}{\omega(1-\tau)}a_2 b_2$. As $a_1 \sim \omega^{\frac{2}{3}}$ we know that the left-hand side will be small and $a_1$ and $b_1$ solve the system if the left-hand side equals 0. Writing $a_2 = a_1 + \tilde{a}_2$ and $b_2 = b_1 + \tilde{b}_2$ this gives:

$$\frac{d}{d\tau}a_1 = 1 - \frac{2}{\omega(1-\tau)}(a_1 + \tilde{a}_2)(b_1 + \tilde{b}_2) \approx \frac{2}{\omega(1-\tau)}(a_1\tilde{b}_2 + b_1\tilde{a}_2) . \tag{S40}$$

From dimensional analysis it follows that the correction terms $\tilde{a}_2$ and $\tilde{b}_2$ must scale like $\tilde{a}_2 \sim \omega^{\frac{4}{3}}$ and $\tilde{b}_2 \sim \omega$ and are hence much smaller than the first order approximations $a_1$ and $b_1$. Higher order corrections will give even smaller contributions showing that if $\frac{d}{d\tau}a_1 \ll 1$, $a_1$ is indeed a very good approximation.

In the limit $\tau \to 0$, however, the expression for $a(\tau)$ in Eq. (S39) diverges and consistency is violated. Hence, the obtained solution is valid only for sufficiently large $\tau$.

We fix some small $\epsilon > 0$ such that the approximation can be assumed to be sufficiently good if $\frac{d}{dt}a < \epsilon$. Furthermore, we define $\tau_\epsilon$ such that $\frac{d}{d\tau}a < \epsilon$ for all $\tau > \tau_\epsilon$. Using Eq. (S39) we can write this as $\frac{d}{d\tau}h < \epsilon\eta^{\frac{1}{3}}/\omega^{\frac{2}{3}}$ for all $\tau > \tau_\epsilon$, where the left-hand side, $\frac{d}{d\tau}h$, depends only on $\tau$. Hence, by decreasing $\omega$ we can make $\tau_\epsilon$ arbitrarily small: $\lim_{\omega\to 0}\tau_\epsilon = 0$. In order to calculate $g(\omega, \eta)$ the integral in Eq. (S36) can be separated in a domain where the approximation $a(\tau)$ is accurate and a domain where the correct solution $\tilde{a}(\tau)$ deviates strongly from $a(\tau)$:

$$g(\omega,\eta) = \frac{1}{\omega}\int_0^{\tau_\epsilon} \frac{\tilde{a}(\tau)}{1-\tau}d\tau + \frac{1}{\omega}\int_{\tau_\epsilon}^{1} \frac{a(\tau)}{1-\tau}d\tau. \tag{S41}$$

We see from Eq. (S35a) that $\frac{d}{d\tau}\tilde{a} = 1$ describes an upper bound to $\tilde{a}$ showing that $\tilde{a}(\tau) \leq \tau$. Therefore we can bound the contribution of the first integral as $\int_0^{\tau_\epsilon} \frac{\tilde{a}(\tau)}{1-\tau}d\tau \leq \int_0^{\tau_\epsilon} \frac{\tau}{1-\tau_\epsilon}d\tau = \frac{1}{2}\frac{\tau_\epsilon^2}{1-\tau_\epsilon}$. Because this upper bound for the integral goes to 0 if $\omega$ and hence $\tau_\epsilon$ become small the first integral will become negligible against the second one. Asymptotically, we therefore only need to consider the second integral with the solution for $a(\tau)$ as given by Eq. (S39):

$$\begin{aligned} g(\omega,\eta) &= (\omega\eta)^{-\frac{1}{3}}\int_0^1 (6t-3t^2)^{-\frac{1}{3}}dt = (\omega\eta)^{-\frac{1}{3}}\int_0^3 \frac{dz}{6z^{\frac{1}{3}}\sqrt{1-\frac{z}{3}}} = \\ &= \frac{3^{\frac{2}{3}}\sqrt{\pi}\,\Gamma(\frac{2}{3})}{6\,\Gamma(\frac{7}{6})}(\omega\eta)^{-\frac{1}{3}} \approx 0.8969\cdot(\omega\eta)^{-\frac{1}{3}}\ , \end{aligned} \tag{S42}$$

where we used the substitution $t = 1 - \sqrt{1 - z/3}$ and $\Gamma(x)$ is the (Euler) Gamma function. So, in the limit of small $\omega$, $g$ scales with $\omega$ and $\eta$ with identical exponent $-\frac{1}{3}$. This contrasts the dimerization scenario where $g$ as well as $A$ and $B$ depend only on $\eta$ and are independent of $\omega$ (cf. Eq. (S32), (S33) and (S34)).

**Numerical analysis and the threshold values for the rate constants** In order to confirm the results of the last two paragraphs and to see how $g(\omega, \eta)$ behaves in the intermediate regime where $\omega$ and $\eta$ are of the same order of magnitude we also investigate the function $g(\omega, \eta)$ numerically. For that purpose we numerically integrate the ODE-system for $A(t)$ and $B(t)$ in Eq. (S26) for different values of $\omega$ and $\eta$ with a semi-implicit method. Subsequently, we integrate the solution $A(t)$ using an adaptive recursive Simpson's rule. Plotting $g$ in dependence of $\omega$ for fixed $\eta$ on a double-logarithmic scale reveals a rather simple bipartite form of $g$, see Fig. S1a:

$$g(\omega,\eta) = \begin{cases} g_1(\eta)\omega^{-\frac{1}{3}} & \omega \ll 1 \\ g_2(\eta) & \omega \gg 1. \end{cases} \tag{S43}$$

The transition between these two regimes is rather sharp so that $g$ is best described in a piecewise fashion

$$g(\omega, \eta) = \max\left(g_1(\eta)\omega^{-\frac{1}{3}}, g_2(\eta)\right). \tag{S44}$$

Next, we plot the coefficients $g_1(\eta)$ and $g_2(\eta)$ against $\eta$. Here we find that $g_1(\eta) = a\eta^{-\frac{1}{3}}$ with $a = \text{const} \approx 0.90$ and $g_2(\eta)$ is again bipartite with a sharp kink in between (Fig. S1b):

$$g_2(\eta) = \min\left(b\eta^{-\frac{1}{2}}, b'\eta^{-0.85}\right), \tag{S45}$$

where $b \approx 1.11$ and $b' \approx 1.37$. The transition between both regimes is at $\eta \approx 1.82$. The second regime is not relevant for self-assembly since it refers to both large $\omega$ and large $\eta$, hence the travelled distance $2g$ is too small to give finite yield in this regime. Therefore, we discard the second regime and obtain as final result

$$g(\omega, \eta) = \max\left(a(\eta\omega)^{-\frac{1}{3}}, b\eta^{-\frac{1}{2}}\right), \tag{S46}$$

with $a \approx 0.90$ and $b \approx 1.11$. This confirms perfectly the exponents as well as the coefficients found in the last two paragraphs. It is, however, surprising that there is such a sharp transition between both regimes, which allows to define $g(\omega, \eta)$ in a piecewise fashion. This behavior must be the result of a series of lower oder terms in $g(\omega, \eta)$ which are unimportant in the limits $\omega \ll \eta$ and $\eta \ll \omega$ but cause the sharp transition when $\omega$ and $\eta$ are of the same order of magnitude.

Finally, we return to our original task of finding the threshold values of the activation and dimerization rate for the onset of yield. Using our result for $g(\omega, \eta)$ in Eq. (S23) we find as necessary and sufficient condition to obtain finite yield in the deterministic system:

$$2\max\left(a(\eta\omega)^{-\frac{1}{3}}, b\eta^{-\frac{1}{2}}\right) \geq L - \sqrt{L}\,. \tag{S47}$$

Alternatively, we can state this result as two separate conditions out of which at least one must be fulfilled to obtain finite yield:

$$2a(\eta\omega)^{-\frac{1}{3}} \geq L - \sqrt{L} \qquad \Rightarrow \qquad \alpha < \alpha_{\text{th}} := P_\alpha \frac{\nu}{\mu} \frac{\nu C}{(L - \sqrt{L})^3} \tag{S48}$$

$$\text{or} \qquad 2b\eta^{-\frac{1}{2}} \geq L - \sqrt{L} \qquad \Rightarrow \qquad \mu < \mu_{\text{th}} := P_\mu \frac{\nu}{(L - \sqrt{L})^2} \tag{S49}$$

where $P_\alpha = 8a^3 \approx 5.77$ and $P_\mu = 4b^2 \approx 4.93$. This verifies Eq. (1) in the main text.

#### S4 Influence of the implementation of sub-nucleation reactions

In the main text we focused our discussion on irreversible binding $L_{nuc} = 2$. In this section we investigate the effect of different implementations of the sub-nucleation reactions.

In general, perfect yield is trivially achieved if the complete ring is the only stable structure. However, yield can be maximal already for smaller nucleation sizes $L_{nuc}$ depending on the explicit decay rate $\delta$. In the deterministic limit without the dimerization and activation mechanisms ($\mu = \nu$, $\alpha \to \infty$ ) a rapid transition from zero yield to perfect yield occurs in dependence of the critical nucleation size (see Fig. S2). The threshold value in this case is approximately half the ring size and is weakly affected by the decay rate $\delta$. In order to obtain finite yield for small nucleation sizes, an extremely high decay rate would be necessary. Hence, maximizing the yield solely by increasing the nucleation size is not very feasible.

In our model, the subcritical reaction rates $\mu_i$ may take different values. Here, we want to restrict our discussion to two scenarios. First, all rates have an identical value $\mu_i = \mu$ and second, the rates increase linearly up to the super-nucleation reaction rate: $\mu_i = \mu + (\nu - \mu) \frac{i-1}{L_{nuc}-1}$.

In the deterministic limit, both implementations show the same qualitative behavior as the dimerization mechanism with $L_{nuc} = 2$ in the main text (see Fig. S3). The only relevant aspect for the final yield is the extend to which nucleation is slowed down in total. In the constant scenario all reaction steps contribute equally. As a results there is a strong dependence on the number of such reaction steps, i.e. on the critical nucleation size. If however, the reaction rates increase linearly with the size of the polymers the dimerzation rate dominates. Only in the case $\mu \ll \nu$ finite yield is observed at all. In this limit the dimerization rate is much smaller than the subsequent growth rates. The explicit form of the different $\mu_i$ is not of major importance for the yield. The total slowdown of nucleation is the central feature. Structure decay does not play any role for intermediate nucleation sizes.

The last question we want to address is how the combination of activation and dimerization mechanism and the corresponding non-monotonic behavior is affected by the nucleation size. Again, we compare constant sub-nucleation growth with a linearly increasing growth rate (see Fig. S4). In the deterministic regime both implementations behave qualitatively similar as the dimerization mechanism discussed in the main text. However, in both cases the stochastic yield catastrophe is less pronounced. For the constant growth rates a saturation of the maximal yield is observed for sufficiently low $\mu$. If the profile is linear this effect is weaker as compared to the constant case and a dependency on the explicit value of $\mu$ is still observed. The saturation value is not reached for these reactions rates.

Taking all our results for the sub-nucleation behavior together we draw the following conclusions: First, structure decay by itself it not very efficient in order to maximize yield. Second, the explicit choice of the sub-nucleation rates is of minor importance for the qualitative behavior. The system behaves similarly to the case $L_{nuc} = 2$. Third, larger nucleation sizes mitigate the stochastic yield catastrophe in general.

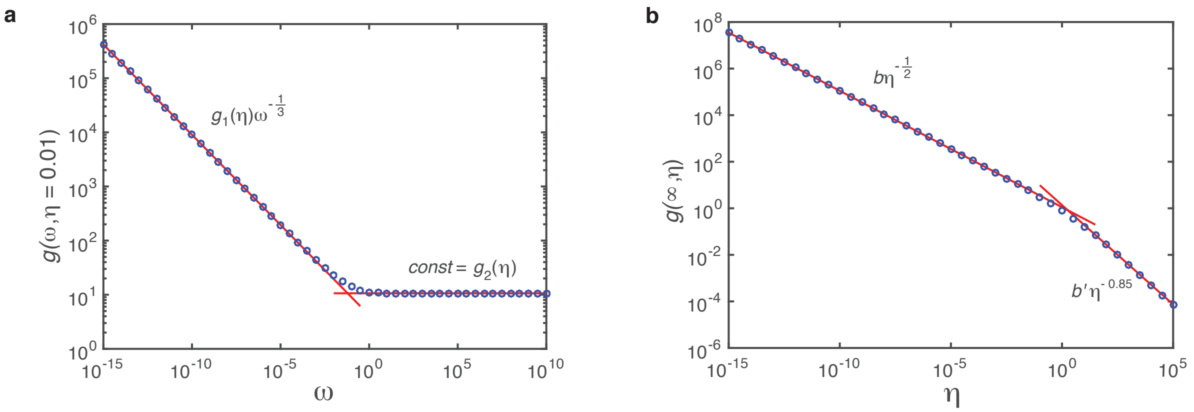


Figure S1: Fit of $g(\omega,\eta)$ on log-log scale. The function $g(\omega,\eta) = \int_0^\infty A_{\omega,\eta}(t)dt$ describes (half) the travelled distance of the profile of the polymer size distribution in dependence of $\omega = \frac{\alpha}{\nu C}$ and $\eta = \frac{\mu}{\nu}$. Marker points show solutions for $g(\omega,\eta)$ as obtained numerically from integration of Eq.(17). Red lines are linear fits on log-log scale. In **a)** we plot $g(\omega,\eta)$ for fixed $\eta$ (here exemplarily for $\eta = 0.01$) over 25 orders of magnitude in $\omega$ and find a markedly bipartite behavior: For small $\omega$ the dependence on $\omega$ is perfectly matched by a power law with exponent $-\frac{1}{3}$ and $\eta$-dependent coefficient $g_1(\eta)$, whereas for large $\omega$ it is a constant $g_2(\eta)$. **b)** Plotting $g_2(\eta) = g(\omega=\infty,\eta)$ in dependence of $\eta$ reveals again strictly bipartite behavior. Here, however, only the brach for small $\eta$ is realistically relevant. With the coefficient $g_1(\eta)$ that can be determined in a similar way this leads to the final form of $g(\omega,\eta)$ as given by Eq. (S46).

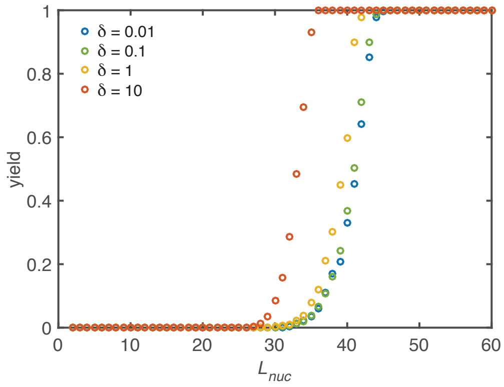


Figure S2: Yield maximization due to increased nucleation size. Without activation and dimerization mechanism $(\alpha \to \infty, \mu = \nu)$ the yield can still be optimized by increasing the critical nucleation size $L_{nuc}$. However, a significant improvement is only achieved for critical sizes larger than half the ring size. Above, a rapid transition to perfect yield takes place. Below no effect is observed at all. Increasing $\delta$ shifts the onset of yield to slightly smaller critical nucleation sizes. Other parameters: $L = 60$, $N = 10000$.

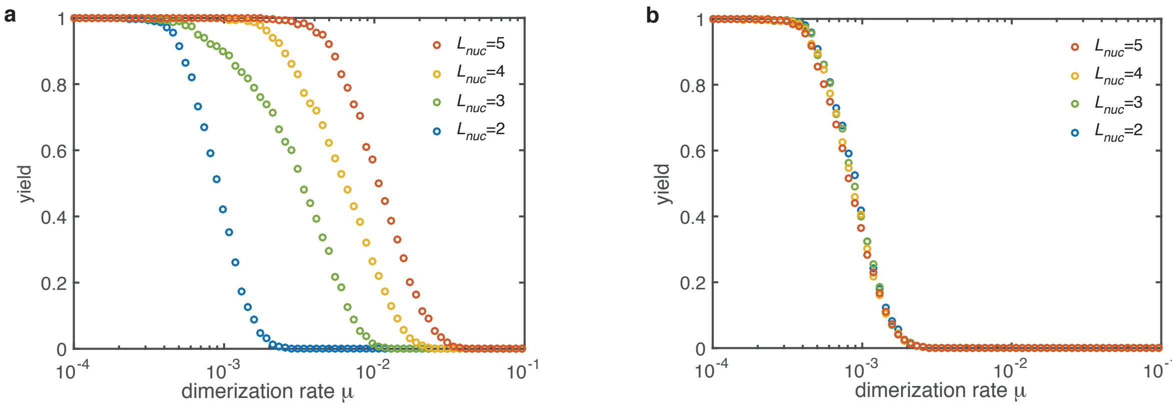


Figure S3: Yield for the dimerization mechanism ($\alpha \to \infty$) with different nucleation sizes (colors). **a** If all sub-nucleation growth rates are identical ($\mu_i = \mu$) increasing the nucleation size increases the threshold value $\mu_{th}$. The slow down of nucleation due to the individual sub-nucleation steps in total determines the yield. **b** If the sub-nucleation growth rates increase linear $\left(\mu_i = \mu + (\nu - \mu)\frac{i-1}{L_{nuc}-1}\right)$ no dependence on the nucleation size is observed. The dimerization rate $\mu_1 = \mu$ (which is the most limiting step) dominates entirely. Other parameters: $L = 60$, $N = 10000$, $\delta = 1$.

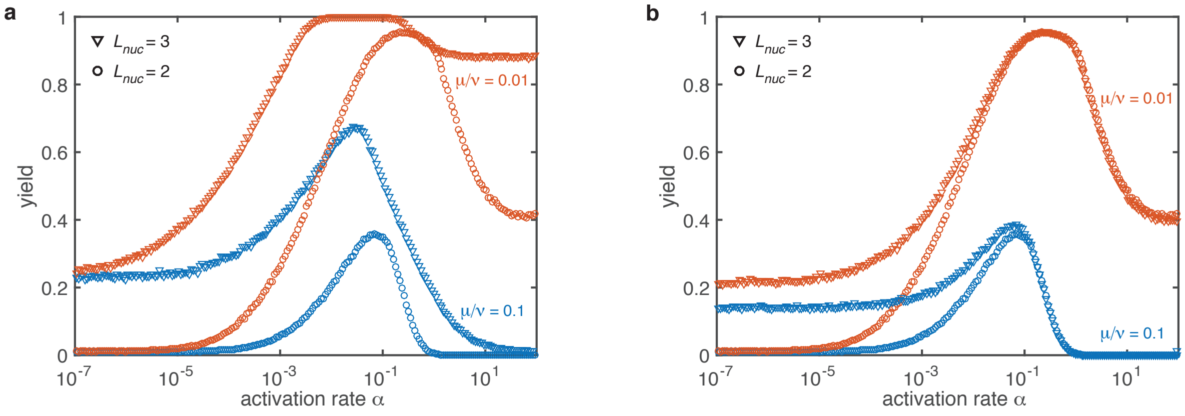


Figure S4: Combined mechanisms for different nucleation sizes (symbols) and dimerization rates (color). **a** If the sub-nucleation growth rates are identical ($\mu_i = \mu$) The stochastic yield catastrophe is weakened but still has a drastic impact. The qualitative behavior remains unchanged. **b** For a linearly increasing sub-nucleation growth rate $\left(\mu_i = \mu + (\nu - \mu)\frac{i-1}{L_{nuc}-1}\right)$ in the deterministic regime no changes are observed at all. The effect of the stochastic yield catastrophe is less pronounced. This improvement is mainly caused by structure decay which mitigates stochastic fluctuations. However, a slight dependency of the saturation value on the rate $\mu$ is observed. Other parameters: $L = 60$, $S = L$, $N = 100$, $\delta = 0.1$.
